# Limited oxygen availability in standard cell culture alters metabolism and function in terminally-differentiated cells

**DOI:** 10.1101/2022.11.29.516437

**Authors:** Joycelyn Tan, Sam Virtue, Dougall M. Norris, Olivia J. Conway, Ming Yang, Christopher Gribben, Fatima Lugtu, Ioannis Kamzolas, James R. Krycer, Richard J. Mills, Conceição Pereira, Martin Dale, Amber S. Shun-Shion, Harry J. M. Baird, James A. Horscroft, Alice P. Sowton, Marcella Ma, Stefania Carobbio, Evangelia Petsalaki, Andrew J. Murray, David C. Gershlick, James E. Hudson, Ludovic Vallier, Kelsey H Fisher-Wellman, Christian Frezza, Antonio Vidal-Puig, Daniel J. Fazakerley

**Affiliations:** Metabolic Research Laboratories, Wellcome-Medical Research Council Institute of Metabolic Science, University of Cambridge, Cambridge CB2 0QQ, UK; MRC Cancer Unit, University of Cambridge, Cambridge Biomedical Campus, Cambridge CB2 0XZ, UK; CECAD Research Center, Faculty of Medicine, University Hospital Cologne, Cologne 50931, Germany; Wellcome-MRC Cambridge Stem Cell Institute, University of Cambridge, Cambridge CB2 0AW, UK; European Molecular Biology Laboratory, European Bioinformatics Institute, Wellcome Genome Campus, Hinxton CB10 1SD, UK; QIMR Berghofer Medical Research Institute, Brisbane, Queensland 4006, Australia.; Faculty of Health, School of Biomedical Sciences, Queensland University of Technology, Brisbane, Queensland 4000, Australia; Cambridge Institute for Medical Research, University of Cambridge, Cambridge CB2 0XY, UK; Department of Physiology, Development and Neuroscience, University of Cambridge, Cambridge CB2 3EL, UK; Centro de Investigacion Principe Felipe, Valencia 46012, Spain; Faculty of Medicine, School of Biomedical Sciences, The University of Queensland, Brisbane, Queensland 4072, Australia; Department of Physiology, Brody School of Medicine, East Carolina University, Greenville, North Carolina 27834, USA; East Carolina Diabetes and Obesity Institute, East Carolina University, Greenville, North Carolina 27834, USA; UNC Lineberger Comprehensive Cancer Center, University of North Carolina at Chapel Hill School of Medicine, Chapel Hill, North Carolina 27599, USA

**Keywords:** Cell culture, oxygen tension, oxygen diffusion, hypoxia, HIF, glucose metabolism, *de novo* lipogenesis, adipocytes, hPSC

## Abstract

Cell culture is generally considered to be hyperoxic. However, the importance of cellular oxygen consumption is often underappreciated, with rates of oxygen consumption often sufficient to cause hypoxia at cell monolayers. We initially focused on cultured adipocytes as a terminally differentiated cell-type with substantial oxygen consumption rates to support diverse cellular functions. Under standard conditions, cultured adipocytes are hypoxic and highly glycolytic. Increasing oxygen diverted glucose flux toward mitochondria and resulted in thousands of gene expression changes that pointed toward alleviated physiological transcriptional responses to hypoxia. Phenotypically, providing more oxygen increased adipokine secretion and rendered adipocytes more sensitive to insulin and lipolytic stimuli. The functional benefits of increasing pericellular oxygen were transferable to other cellular systems including hPSC-derived hepatocytes and cardiac organoids. Our findings suggest that oxygen is limiting in many terminally-differentiated cell culture systems, and that controlling oxygen availability can improve the quality and translatability of cell models.

## INTRODUCTION

Over the past two decades, substantive focus has been placed on improving translation between mice and humans^1–3^. However, until recently, attempts to improve translation between cell culture and *in vivo* settings, be they mouse or human, has not received as much attention. In the past 5-6 years there have been efforts to improve the fidelity of cell culture models principally by altering the nutrient composition of cell culture media to be more physiological^4, 5^. However, the importance of oxygen tension in cell culture has not received the same attention.

Cell culture is usually viewed as hyperoxic^6^, due to incubators containing approximately 18% oxygen. As a result, most work studying oxygen tension in cell culture focuses on reducing atmospheric oxygen concentrations to 5% or even 1%. However, the notion that cell culture is hyperoxic relies on an assumption that oxygen delivery through the medium exceeds cellular demand. First, the depth of medium in which cells are cultured is not trivial. Under “standard culture conditions” (e.g. 1 mL of medium per well of a 12-well plate) the depth of medium varies between 2.4 and 2.9 mm due to the meniscus effect. Most mammalian cells range between 40 and 100 µM in diameter. This means that oxygen must diffuse through a distance equivalent to between 27 and 58 cell diameters to get to a cell monolayer. Most cells *in vivo* are directly subtended by capillaries which deliver oxygen over significantly shorter diffusion distances (microns) than the medium column (millimetres). Thus, standard culture conditions pose a significant barrier to oxygen delivery. Second, cultured cells have a large range of oxygen consumption rates^7^. If these exceed the rate of oxygen delivery from the air-medium interface, cells will experience hypoxia at the cell monolayer. While the concept that cellular respiration can outstrip oxygen delivery in cell culture is known^8–17^, the functional consequences of hypoxia under standard culture conditions on cell metabolism, gene expression and function has not been addressed.

Many cultured cell lines exhibit high rates of glycolytic metabolism generating lactate from glucose to meet their ATP demands^18, 19^. This glycolytic metabolism is often attributed to an intrinsic metabolic rewiring of cultured cells, similar to the aerobic glycolysis/Warburg effect observed in cancer cells^20^. Indeed, a high rate of glycolysis is a key metabolic adaptation in proliferating cells, suggesting that this may be a necessary cell-intrinsic adaptation to support proliferation^21–24^. However, this reasoning does not hold for the high rates of glycolysis observed in many terminally-differentiated non-proliferative cells^9, 11, 18^. Non-cell-intrinsic causes for high rates of lactate production by cell lines have been previously investigated.

One suggestion to explain why glycolysis appears almost ubiquitous to cell culture is that unphysiological media compositions promote glycolysis, also referred to as the Crabtree effect^25^. However, an arguably simpler explanation for the high glycolytic rates of many cultured cells is that oxygen is limiting; with cells responding physiologically to a hypoxia by switching to anaerobic metabolism. While data regarding the glucose and lactate metabolism by many terminally-differentiated cell types is consistent with the hypothesis that these cell types are hypoxic, it remains to be formally proven and, perhaps more importantly, the impact on cellular phenotypes of increasing oxygen supply to cells is largely unexplored.

Here, we provide experimental evidence that multiple terminally-differentiated metabolic cell lines are hypoxic under standard culture conditions, and explore the consequences of limited oxygen availability on cellular metabolism and phenotypes. We show that the high glycolytic activity of several cultured cell lines is a physiological response to hypoxia, resulting from limitations in oxygen diffusion through the medium column. We demonstrate that increasing oxygen availability reprogrammed cells away from glycolytic and toward oxidative metabolism, increased pyruvate dehydrogenase (PDH) flux, tricarboxylic acid (TCA) cycle activity and oxygen consumption, promoted lipid anabolism, destabilised hypoxia-inducible factor (HIF) 1α protein, and induced over 3000 transcriptional changes within 16 h. Notably, these changes occurred in multiple cell lines and resulted in improved cellular functions. Our findings highlight that understanding and controlling oxygen availability in cell culture models is important for the development of better *in vitro* models.

## RESULTS

### Increasing pericellular oxygen tension decreases lactate production and increases *de novo* lipogenesis in 3T3-L1 adipocytes

To study the effects of oxygen tension in cell culture, we selected the 3T3-L1 adipocyte cell line as our primary model. 3T3-L1 adipocytes are generated from a fibroblast precursor cell line^26^ and are a valuable model used to study adipocyte differentiation^27–29^, lipid metabolism^30^, and hormonal responses^31^. Under standard culture conditions (e.g., 1 mL DMEM/10% FCS per well in 12-well plates) (Figure S1A), 3T3-L1 adipocytes grow as a monolayer and produce high levels of lactate (Figure 1A)^18^. Given the high oxygen consumption rate (OCR) (200 fmol/mm^2^/s) of 3T3-L1 adipocytes (Figures 1B and 1C), we hypothesised that their glycolytic phenotype may not be innate, but a consequence of limited oxygen availability. Indeed, *in vivo* estimates from human subcutaneous adipose tissue across fasting and postprandial periods place adipose tissue lactate production at 15-30% of glucose^18, 32^, which is lower than observed for cultured 3T3-L1s (close to 50%). Fick’s first law of gas diffusion (Figure 1D) states that oxygen diffusion rates are proportionate to medium depth. To support an OCR of 200 fmol/mm^2^/s we calculated that medium depth could not exceed 2.43 mm (e.g., a 5 mm medium depth would limit OCR to just 100 fmol/mm^2^/s). Medium height is not constant in cell culture wells due to the meniscus. The minimum depth of medium in a 12-well plate containing 1 mL of medium was measured at ∼2.4 mm (Figures 1D and S1A). Based on the OCR of 3T3-L1s, Fick’s law predicts that oxygen concentrations at the cell monolayer would be virtually 0 (Figure 1D). Indeed, under standard culture conditions (100 µL per well in a 96-well plate), the oxygen concentration measured 0.55 mm above the cell monolayer (the minimum measurable depth by the Resipher system) was ∼15 µM (11 mmHg) (Figures 1E and S1A), well below atmospheric oxygen (181 µM; 139 mmHg). To increase oxygen tension at the cell monolayer, we lowered medium volumes, reducing the distance from the air-medium interface to the cell monolayer (Figure 1D). As expected, lowering medium volumes increased pericellular oxygen levels (Figure S1B), with oxygen concentrations 0.55 mm above the cell monolayer reaching between 38–73 µM (29–56 mmHg) for cells cultured in 50 µL or 33 µL medium in a 96-well plate, respectively (Figure 1E). These values were more similar to those measured in mouse (∼60 mmHg)^33^ and human (40.5 – 73.8 mmHg)^34^ adipose tissue, than cells grown under standard conditions. Therefore, lowering medium volumes provided a simple experimental system to assess the impact of oxygen tension on 3T3-L1 adipocyte metabolism^12^.

**Figure 1.**
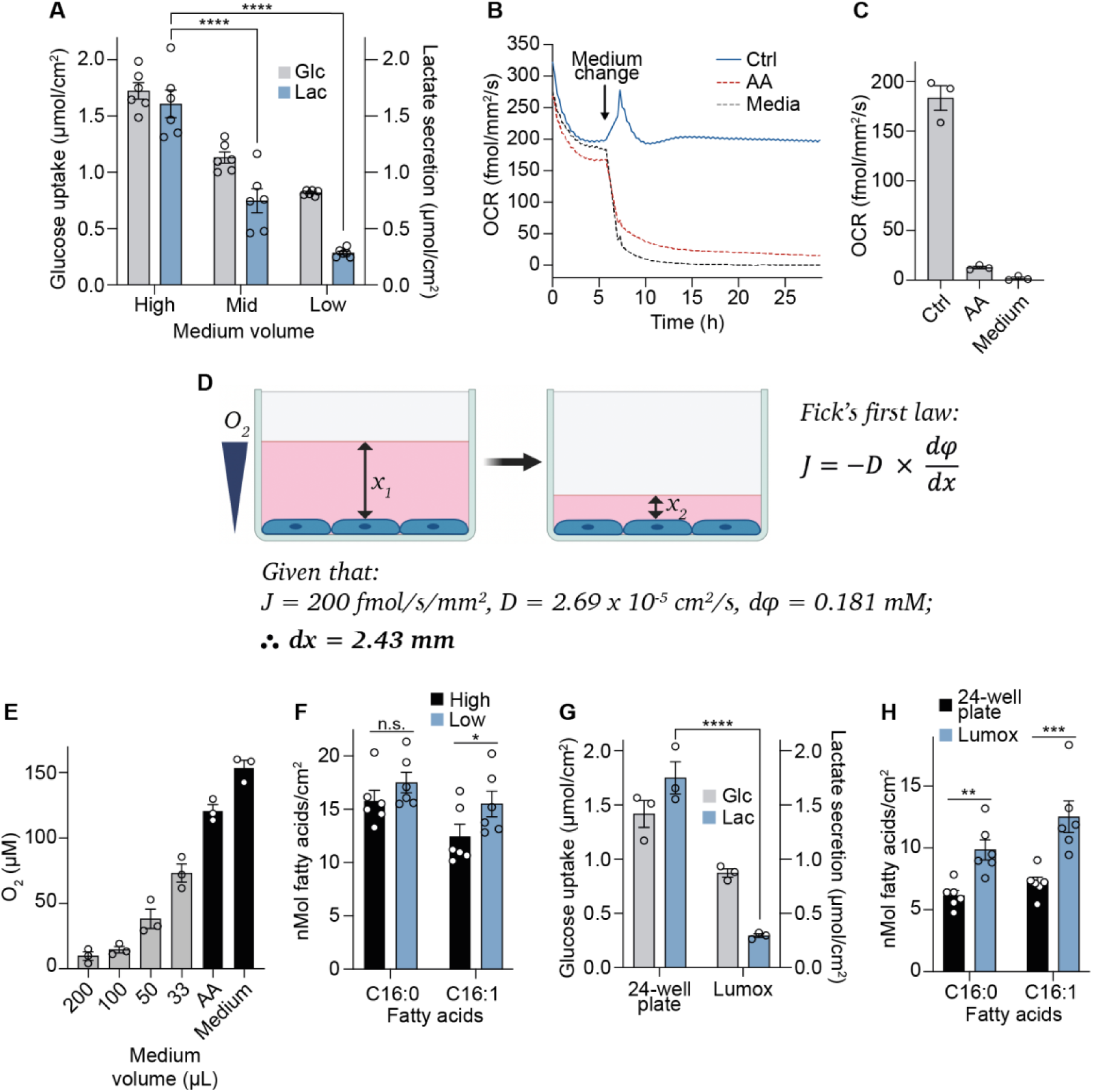
Increasing pericellular oxygen tension decreases lactate production and increases *de novo* lipogenesis in 3T3-L1 adipocytes. A) Extracellular medium glucose and lactate measurements 16 h after medium volume change. High = 1 mL, Mid = 0.67 mL, Low = 0.33 mL (in a 12-well plate) from hereafter unless stated otherwise. (n = 6 biological replicates). (B) 3T3-L1 adipocyte oxygen consumption rate (OCR) under control (100 µL), mitochondrial complex III inhibitor antimycin A (AA) treatment, or cell-free/medium only conditions (96-well plate). Baseline OCR was measured for 6 h before starting AA treatment and removing cells from medium-only wells (representative of n = 3 biological replicates). (C) OCR measurements taken after 20 h from (B) (n = 3 biological replicates). (D) Schematic representation of experimental setup and Fick’s first law of diffusion, which states that the rate of diffusion (*J*) is proportional to the concentration gradient (φ) and is inversely proportional to the diffusion length (*x*), where *D* is the diffusion coefficient. (E) Pericellular oxygen concentration at different medium volumes in 96-well plates (n = 3 biological replicates). (F) *De novo* lipogenesis (DNL) of palmitate (C16:0) and palmitoleate (C16:1) during 16 h of culture in high or low medium volumes in 12-well plates (n = 6 biological replicates). (G) Extracellular medium glucose and lactate measurements in 24-well or Lumox plates after 16 h culture in high or low medium volumes (n = 3 biological replicates). (H) DNL of palmitate and palmitoleate after 16 h in 24-well or gas-permeable Lumox plates (n = 6 biological replicates). Data are represented as mean ± SEM. \**p* < 0.05, \*\**p* < 0.01, \*\*\**p* < 0.001, \*\*\*\**p* < 0.0001 by two-way ANOVA with Šidák correction for multiple comparisons (A, E, F, G, and H). See also Figure S1.

Under low oxygen tension, glycolysis switches from being primarily aerobic, with CO_2_ as the end point, to becoming increasingly anaerobic, with lactate as the end point. We initially measured glucose consumption and lactate production. Switching cells cultured in 12-well plates from 1 mL (high) to 0.67 mL (mid) or 0.33 mL (low) medium lowered glucose uptake and caused an even more drastic reduction in lactate production (Figure 1A). Notably, the extent of lactate production from glucose in 3T3-L1 adipocytes cultured at a higher oxygen tension (17% lactate from glucose) is more similar to the *in vivo* values of 15-30% in human subcutaneous abdominal adipose tissue^32^. This change was not driven by depletion of glucose, which remained >12 mM under all medium volumes after 16 h (Figure S1C). Instead, our results indicated a considerable reduction in anaerobic glycolysis under conditions of greater oxygen availability (Figure 1A). Importantly, the reduction in the lactate-to-glucose ratio in low medium conditions was independent of starting medium glucose concentrations (Figure S1D), suggesting that the effect of increasing oxygen on glucose metabolism was not an artefact of culturing cells in non-physiological medium glucose^25^.

Critically, despite a 50% reduction in glucose uptake, cells cultured in lower medium volumes (0.33 mL *versus* 1 mL) increased *de novo* lipogenesis (DNL) (Figure 1F), indicating the cells were substantially more metabolically efficient; able to produce sufficient ATP and obtain enough carbon to support DNL from less glucose. The increased efficiency of the cells was consistent with the fact that generation of ATP by converting glucose to lactate only produces 2 molecules of ATP, whereas complete oxidation in mitochondria generates up to 36^35^. To confirm these metabolic responses to low medium conditions were due to a primary change in oxygen availability, we cultured cells in gas-permeable cell culture plates (Lumox). These plates allow oxygen diffusion from the bottom of the well, bypassing the media layer. Similar to the low medium conditions, gas-permeable plates lowered glucose use and lactate production (Figure 1G), and increased DNL (Figure 1H). Overall, these results demonstrate that the glycolytic metabolism of 3T3-L1 cells is not innate, but a response to limited oxygen supply. We next sought to determine how important the impact of increasing oxygen tension was to the metabolism and phenotypes of cells in culture.

### Lowering medium volumes increases mitochondrial glucose oxidation

We next directly measured OCR under different medium volumes. Switching cells from standard (100 µL per well in 96-well plates) to low (33 µL) medium conditions rapidly (within 3-4 h) doubled the oxygen consumption rate from 200 to 400 fmol/mm^2^/s (Figure 2A), showing that standard culture volumes were indeed limiting for oxygen delivery to 3T3-L1 adipocytes. Increased O2 consumption was independent of changes in mitochondrial content as measured by western blotting (Figure S2A), or oxygen consumption in permeabilised cells (Figure S2B). Under standard culture conditions, medium oxygen concentration was ∼15 µM

**Figure 2.**
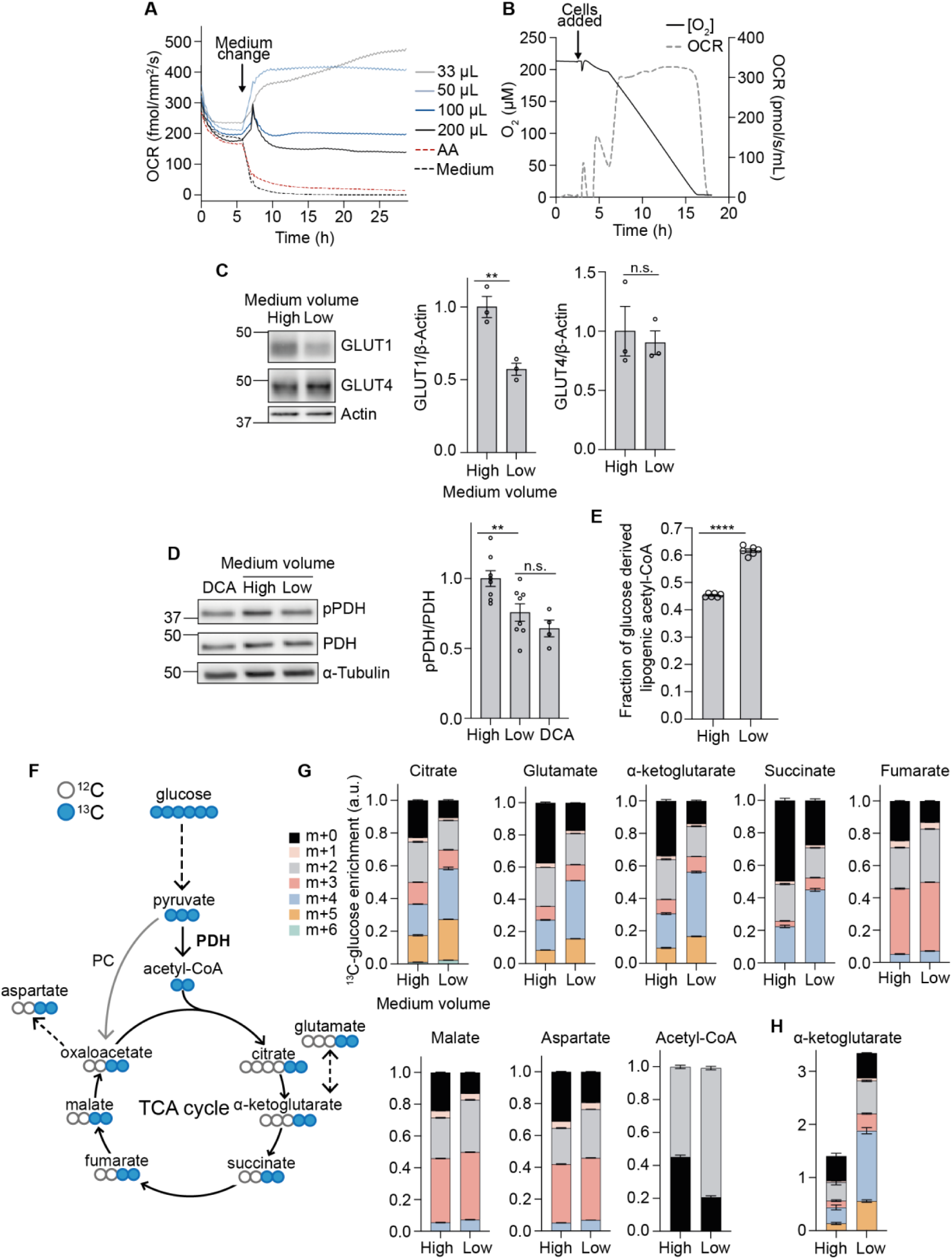
Lowering medium volumes increases mitochondrial glucose oxidation. (A) 3T3-L1 adipocyte oxygen consumption rate (OCR) at different medium volumes, antimycin A (AA) treatment, or medium only conditions in 96-well plates. Baseline OCR (at 100 µL medium) was measured for 6 h before altering medium volumes or starting AA and medium only treatments (n = 3 biological replicates). (B) Oroboros measurements showing oxygen concentration and rate of oxygen consumption (OCR) of 3T3-L1 adipocytes in a closed system (n = 3 biological replicates). (C) Western blot and quantification of GLUT1 and GLUT4 after 16 h of medium volume change (n = 3 biological replicates). (D) Western blot and quantification of phospho-pyruvate dehydrogenase (pPDH) and total PDH after 16 h of medium volume change in 12-well plates, with dichloroacetate (DCA) as a positive control (n = 8 biological replicates). (E) Fraction of newly synthesised lipids after 16 h of medium volume change and U^13^C-glucose labelling in 12-well plates (n = 6 biological replicates). (F) Schematic of ^13^C enrichment in TCA cycle metabolites after one cycle of U^13^C-glucose labelling. PC, pyruvate carboxylase. (G) Graphs show the fractional abundance of each isotopologue after 16 h medium volume change (n = 6 biological replicates). (H) Total abundance of α-ketoglutarate after 16 h medium volume change (n = 6 biological replicates). Data are represented as mean ± SEM. n.s., non-significant; \*\**p* < 0.01, \*\*\*\**p* < 0.0001 by paired two-tailed Student’s t-test (C and D). See also Figure S2.

0.55 mm above the cells (Figure 1E). However, in a closed system, 3T3-L1 adipocytes respired maximally down to ∼4 µM oxygen (Figure 2B). Together, these data suggest a near total use of available oxygen (to less than 4 µM) at the cell monolayer. Overall, we demonstrate that 3T3-L1 adipocytes consume all oxygen that diffuses to the monolayer under standard culture conditions, become hypoxic, and perform anaerobic glycolysis to maximise ATP production.

Multiple adaptations to hypoxia are known, including increased glucose uptake for anaerobic glycolysis^36^, inhibition of PDH flux to limit aerobic TCA cycle activity^37^, and activation of the HIF system^38, 39^. Firstly, we determined changes in glucose transport and expression of glucose transporters. We did not see any changes in the protein levels of the insulin-regulated glucose transporter GLUT4. In contrast, the hypoxia-inducible glucose transporter GLUT1 was strongly suppressed by low medium, as was non-GLUT4 dependent glucose uptake (Figures 2C and S2C). Second, PDH phosphorylation was decreased following the switch to low medium volumes (Figure 2D), consistent with decreased negative regulation of PDH by pyruvate dehydrogenase kinases (PDKs) and therefore increased PDH activity^37, 40^. Indeed, direct measurement of PDH flux using mass isotopomer distribution analysis (MIDA) of palmitate^41^ revealed a ∼40% increase in glucose entry to the TCA cycle via PDH when cells were cultured in lower medium volumes (Figure 2E). Consistent with greater PDH activity and glucose entry into the TCA cycle, U^13^C-glucose tracing (Figure 2F) revealed that transitioning cells to low medium (0.33 mL in 12-well plates) for either 4 h (Figure S2E) or 16 h (Figure 2G) increased ^13^C-labelling of all TCA cycle intermediates (Table S1). Additionally, enrichment of m+3 isotopologues for fumarate and malate suggested increased pyruvate carboxylase activity, which allows for TCA anaplerosis by replenishing oxaloacetate. In total, the abundance of 45 metabolites was changed by culturing cells in low medium for 16 h, indicating comprehensive metabolic rewiring of cells beyond just the TCA cycle (Figure S2G). Of note, α-ketoglutarate (ɑ-KG) was the most upregulated TCA metabolite under low medium conditions (Figure S2F), with all the additional ɑ-KG containing glucose-derived carbon (Figure 2H). These results demonstrate that increasing oxygen tension can rewire multiple branches of cell metabolism in addition to increased glucose oxidation via the TCA cycle.

### Lowering medium volumes induces a widespread transcriptional response and improves adipocyte function

The increase in α-KG abundance, which governs the activity of α-KG-dependent dioxygenase prolyl hydroxylases^42–44^, alongside changes in cellular oxygen tension and lower GLUT1 protein levels^45^, are all indicative of reduced HIF1α stability and decreased transcriptional activity. HIF1α abundance was drastically decreased when cells were switched to low medium for 16 h (Figure 3A). Despite the stabilisation of HIF1α in high medium under standard culture conditions, adipocytes still mounted a considerable HIF1a stabilisation and lactate response to further hypoxia (5% oxygen; Figures S3A and S3B). The partial reversal of both HIF1α stabilisation and lactate production under low medium conditions at 5% oxygen support our conclusion that low medium affects these parameters through greater oxygen provision (Figures S3A and S3B). In further agreement, expression of HIF1α target genes (*Slc16a3, Pgk1, Pdk1, Car9, Pkm2, Slc2a1*) was also reduced under low medium conditions (Figure 3B) and in cells cultured in gas-permeable culture plates (Lumox) (Figure S3C). These data demonstrate clearly that the 3T3-L1 cell line has a regulatable and functional oxygen-responsive HIF system, which is active under standard culture conditions.

**Figure 3.**
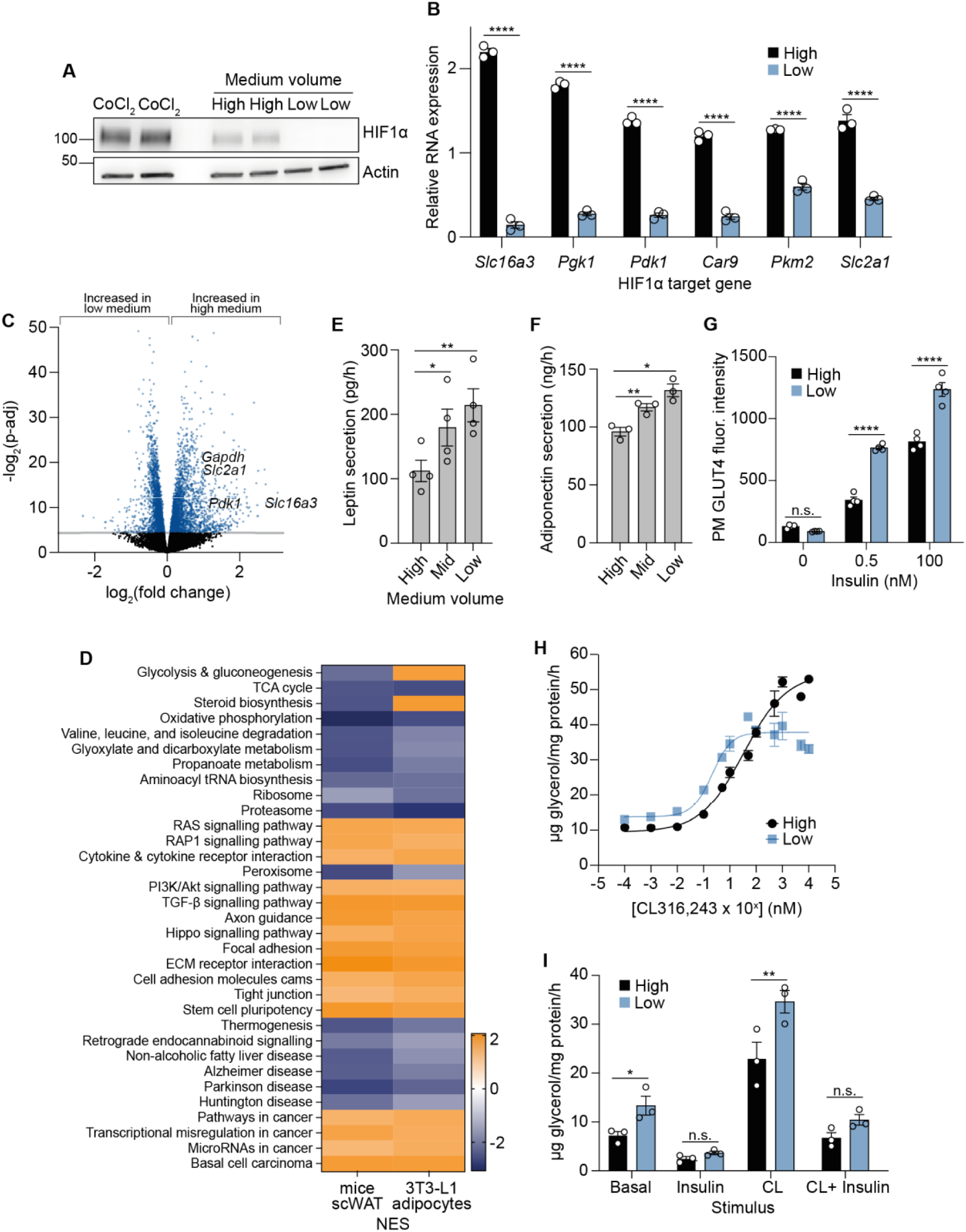
Lowering medium volumes induces a widespread transcriptional response and improves adipocyte function. (A) Western blot of hypoxia-inducible factor (HIF) 1α after 16 h medium volume change in 12-well plates, with 500 µM CoCl2 as positive control (n = 3 biological replicates). (B) Relative RNA expression of HIF1α target genes in 3T3-L1 adipocytes after 16 h medium volume change in 12-well plates (n = 3 biological replicates). (C) Volcano plot of differentially expressed genes after 16 h medium volume change in 12-well plates (n = 6 biological replicates). (D) KEGG pathway analyses of 3T3-L1 adipocytes after 16 h medium volume change in 12-well plates (n = 6 biological replicates) and subcutaneous white adipose tissue (scWAT) from mice kept in 10% or 21% oxygen for 4 weeks (n = 10 biological replicates). Orange (positive NES) represents upregulated KEGG pathways in high medium (3T3-L1 adipocytes) or in 10% oxygen (mice scWAT). Blue (negative NES) represents upregulated KEGG pathways in low medium (3T3-L1 adipocytes) or in 21% oxygen (mice scWAT). NES, normalised enrichment score. (E) Rate of leptin secretion under different medium volume conditions in 12-well plates (n = 4 biological replicates). (F) Rate of adiponectin secretion under different medium volume conditions in 12-well plates (n = 4 biological replicates). (G) Fluorescence intensity of plasma membrane (PM) GLUT4 upon insulin stimulation after 48 h medium volume change in 96-well plates (n = 4 biological replicates). (H) Dose response curve of lipolytic drug, CL316,243 treatment in 12-well plates (n = 7 biological replicates). (I) Rate of lipolysis measured by glycerol release, upon 100 nM insulin or 1 nM CL316,243 stimulation (n = 4 biological replicates). Data are represented as mean ± SEM. \**p* < 0.05, \*\**p* < 0.01, \*\*\**p* < 0.001, \*\*\*\**p* < 0.0001 by one/two-way ANOVA with Šidák correction for multiple comparisons (D, E, F, and H), or by paired Student’s t-tests (B). See also Figure S3.

To fully characterise the transcriptional responses to low medium we performed RNAseq analyses. This revealed extensive and widespread changes in gene expression in cells cultured in low medium, with over 3000 differentially expressed genes. Multiple HIF1α targets were expressed at greater levels in high medium (Figure 3C; Table S2). VIPER transcription factor prediction software predicted 125 transcription factors to either be inhibited or activated (Table S3) by changing medium volumes, including HIF1α (NES 8.8, FDR P-value=1.6 10^-16^). Finally, FGSEA identified 63 altered KEGG pathways (Table S4). Notably, mitochondrial function, TCA cycle activity, and lipogenesis, were all predicted to be increased by low medium (Table S4), consistent with our biochemical analyses (Figures 1 and 2).

Given the widespread transcriptional response to increased oxygen tension in cultured cells, we next compared our *in vitro* findings to *in vivo* adipose tissue. We compared the transcriptional profile of 3T3-L1 adipocytes in low *versus* high medium culture conditions to subcutaneous white adipose tissue (scWAT) from mice exposed to either normoxia (21% O_2_) or hypoxia (10% O_2_, 4 weeks) (Figure S3D, Tables S5 and S6). Both individual gene and pathway analyses revealed a high degree of overlap in gene expression responses of cells cultured in high medium and hypoxic adipose tissue. Specifically, of the 2239 differentially expressed genes in hypoxic mice scWAT, 703 were also changed in the 3T3-L1 adipocytes cultured in high medium (Figure S3E), with a 77% concordance in the direction of change (Fig S3F). Additionally, out of 312 KEGG pathways tested, 63 were significantly altered in 3T3-L1 adipocytes and 61 in scWAT, of which 34 were significant in both (Chi^2^ for overlap p=6*10E^-11^) (Tables S4 and S6). Remarkably, 32 of the 34 common pathways were concordantly regulated (Figure 3D, Tables S4 and S6). Overall, our data suggested that increasing oxygen concentrations relative to “standard” culture conditions drove a similar pattern of transcriptional changes to that seen between normoxic and hypoxic adipose tissue, demonstrating that the changes observed in 3T3-L1 adipocytes were relevant to *in vivo* adipose tissue biology.

The extensive changes in gene expression suggested that increased oxygen availability to cells may have broader implications than altered glucose metabolism. Therefore, we next tested cellular phenotypes relevant to adipocyte biology including adipokine secretion, insulin responsiveness, and lipolysis. We cultured adipocytes in high or low medium volumes for 48 h, during which their metabolic switch to glucose oxidation in low medium was maintained (Figure S3G). At this time point, low medium increased both leptin and adiponectin secretion (Figures 3E and 3F), consistent with gene expression data for *Lepn* and *AdipoQ* (Table S2). Additionally, low medium markedly increased insulin-stimulated GLUT4 translocation at both physiological and supraphysiological insulin concentrations (Figure 3G). Similarly, while total 2-deoxyglucose (2-DG) uptake in the presence of insulin was similar in cells cultured in low or high medium volumes, the treatment of cells with the GLUT4-inhibitor indinavir demonstrated that 2-DG uptake was more GLUT4-dependent in low medium (Figure S2D), despite no changes in total GLUT4 protein (Figure 2C).

While adipokine secretion and insulin-stimulated glucose uptake are important functions of mature adipocytes, the primary role of adipose tissue is the storage and mobilisation of fat. Extended culture in low medium increased the sensitivity of 3T3-L1 adipocytes to the lipolysis-stimulating β3-adrenergic receptor agonist CL316,243 more than 15-fold (EC_50_ 0.21 nM *versus* 3.43 nM) (Figure 3H). Despite higher lipolytic sensitivity, insulin was equally effective at suppressing lipolysis in response to CL316,243 (Figure 3I). Overall, these phenotypic measures suggested that increased oxygen tension improved endocrine, glucose, and lipid metabolic functions in 3T3-L1 adipocytes.

### Lowering medium volumes reduces lactate production and improves functional outcomes in other cell types and organoids

Finally, we extended our analysis to other cell types/tissues. First, we assessed lactate secretion in other post-mitotic cell lines cultured under high or low medium volume conditions. pBAT cells (murine brown adipocytes) and L6 myotubes (rat) responded to lower medium volumes by significantly reducing lactate secretion (Figure 4A). Next, we extended our studies to assess the functional effects of increasing oxygen tension in human pluripotent stem cells (hPSC)-derived hepatocytes and cardiac organoids. Culturing induced pluripotent stem cells (iPSC)-derived hepatocytes in low medium lowered lactate secretion (Figure 4B) and HIF1α target gene expression (Fig S4A), indicating that lowering medium volumes increased oxygen tension. Concomitant with changes in HIF1a activity, hepatocyte differentiation markers *ALB* (albumin) and *SERPINA1* were increased, and the lineage specification marker *HNF4α* was decreased^46, 47^ (Figure 4C). On a protein level, we observed increased CYP3A4 activity and hepatocyte albumin content (Figures 4D and 4E). Overall our results suggested that lower media volumes, which decreased HIF1a activity, may increase the functionality of hepatocytes derived from iPSCs^9, 48^. hPSC-derived cardiac organoid cultures also secreted less lactate when cultured in lower medium volumes (Figures 4F and S4B). Organoids in low medium exhibited greater contractile force (Figure 4G), without changes in contractile rate, or activation and relaxation times. (Figures S4C–E). Our data suggest that manipulating oxygen tension, at least in terminally-differentiated cells that model major metabolic organs, impacts the metabolism and phenotype of multiple different cell types.

**Figure 4.**
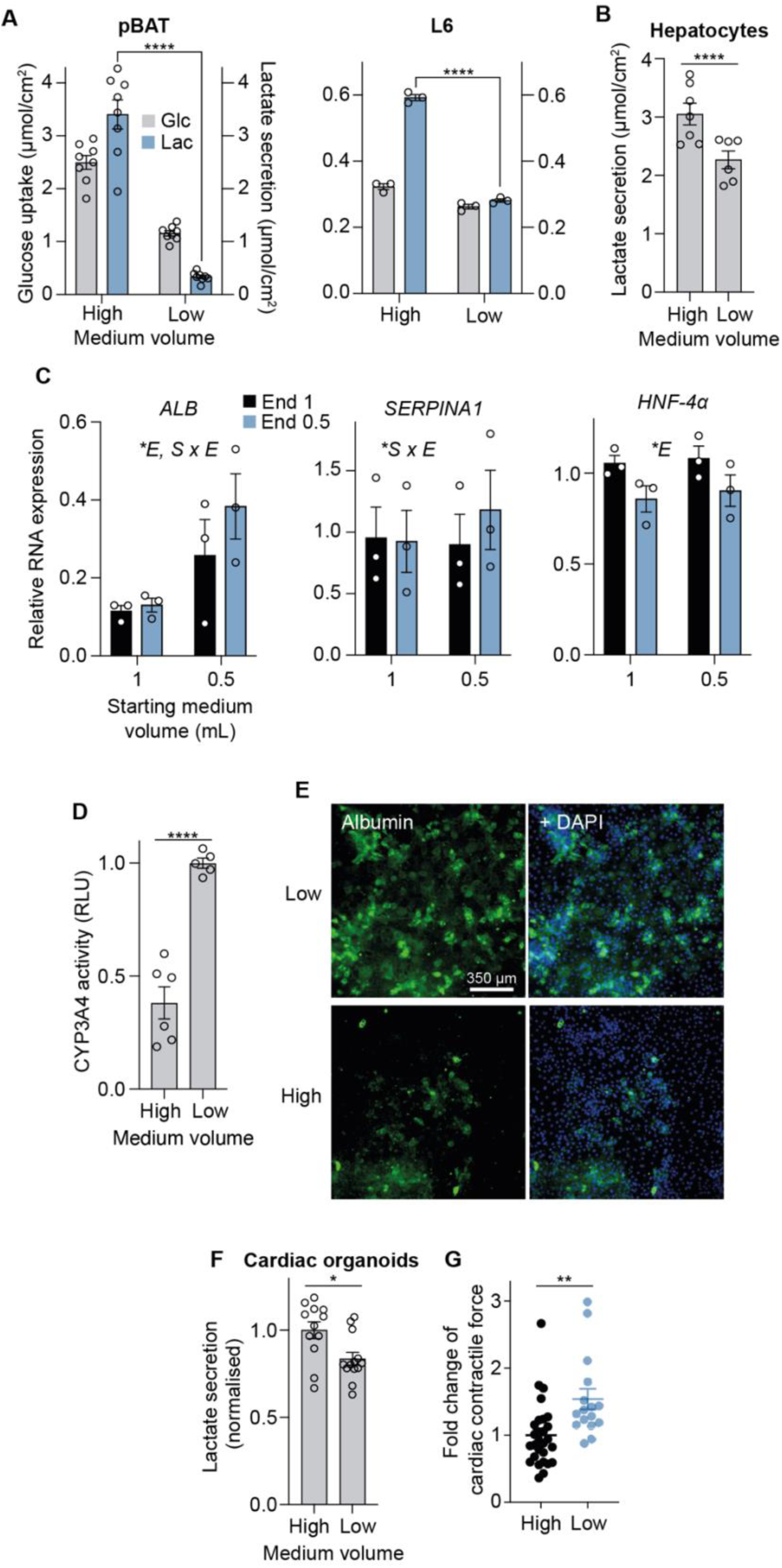
Lowering medium volumes reduces lactate production and improves functional outcomes in other cell types and organoids. (A) Extracellular medium glucose and lactate measurements after 16 h medium volume change in 12-well plates in murine brown adipocytes (pBAT) (n = 8 biological replicates) and L6 myotubes (n = 3 biological replicates). (B) Lactate secretion in iPSC-derived hepatocytes (High = 1 mL, Low = 0.5 mL) after 24 h of medium volume change in 12-well plates (n = 7 biological replicates). (C) Relative RNA expression of hepatocyte differentiation marker genes. Cells were cultured in either 1 mL or 0.5 mL of medium in 12-well plates throughout differentiation, and switched to either 1 mL or 0.5 mL of medium 24 h prior to the experiment (n = 3 biological replicates). \**E* = significance due to end medium volume, \**S* = significance due to starting medium volume, \**S x E* = significance due to interaction between starting and end media volumes. (D) Relative CYP3A4 activity in iPSC-derived hepatocytes after 24 h of medium volume change (n = 6 biological replicates). High = 1 mL, Low = 0.5 mL. (E) Immunofluorescence of albumin in iPSC-derived hepatocytes after 24 h of medium volume change (n = 3 biological replicates). Scale bar = 350 µm. (F) Lactate secretion in cardiac organoids after 48 h of medium volume change in 96 -well plates (normalised) (n = 3-6 technical replicates from n = 3 biological replicates). High = 150 µL, Low = 50 µL. (G) Cardiac contractile force (normalised) (n = 2-17 technical replicates from n = 4 biological replicates). All data are represented as mean ± SEM. \**p* < 0.05, \*\**p* < 0.01, \*\*\**p* < 0.001, \*\*\*\**p* < 0.0001 by two-way ANOVA with Šidák correction for multiple comparisons (A, C, and F) or by paired/unpaired two-tailed Student’s t-test. (B, D, G, and H). See also Figure S4.

## DISCUSSION

Using a comprehensive series of biochemical, multi-omics, and phenotypic analyses, we demonstrate that multiple terminally-differentiated metabolic cell models are hypoxic under standard culture conditions. That oxygen tension can be limiting in standard cell culture has been described and discussed previously^8–17^. However, this is not widely considered, perhaps due to a lack of data on how oxygen deficiencies actually affect cells in culture both metabolically and phenotypically. Our study builds upon this previous work, to provide a comprehensive investigation on the broader impact of oxygen limitations in standard cell culture. Importantly, we show that increasing oxygen availability resulted in significant metabolic and transcriptional changes, and markedly improved cellular function in several cell types that model major metabolic organs – namely adipocytes, hiPSC-derived hepatocytes, and hPSC-derived cardiac organoid cultures. These data provide empirical evidence that limited oxygen has profound effects on cell metabolism and function under standard culture conditions.

There are several possibilities how increased oxygen availability in culture may improve or alter cellular function. The observed impact on cellular phenotypes may be HIF-driven, given the wide range of cellular functions influenced by HIF^38, 49^. For example, we measured increased rates of lipolysis in cells with greater oxygen availability, consistent with previous findings^50^. However, our RNAseq analyses revealed transcriptional changes beyond the HIF pathway (Figure 3), suggesting that additional pathways may connect oxygen availability to cellular behaviour/responses. Indeed, signalling molecules such as reactive oxygen species^51^, or metabolites^52^ may be implicated. For example, in the abundance of a number of metabolites altered upon increased oxygen including lactate, acetyl-CoA, succinate, fumarate, and α-ketoglutarate can affect cellular signalling through post-translational modifications, metabolite receptors, and/or epigenetics^52–54^. Our simple method to increase oxygen to cells, which induces a large shift in cell metabolism, may provide a useful system to further interrogate the link between oxygen tension, metabolites, and cell function.

We used two methods to increase oxygen availability to terminally-differentiated metabolic cell-types; decreasing medium volumes to decrease oxygen diffusion distances, as well as gas-permeable culture plates. Lowering medium volume provides an accessible method to increase oxygen provision which can be easily employed by most laboratories. However, this method has its limitations, such as medium evaporation and nutrient depletion. Other methods to increase cellular oxygen have been employed^7^, but bioreactors or perfusion systems most accurately recapitulate the sophisticated oxygen delivery system of the vasculature since they continuously supply oxygenated medium^55^. However, scalability of and accessibility to this method pose major barriers to widespread use. The field where oxygen tension has perhaps been most carefully considered is in organoid culture. Air−liquid interface culture models have been adopted for many stem-cell and organoid lines, including the respiratory tract^56^, brain^57^, tumour immune microenvironment^58^, and gastrointestinal tissues^59, 60^, with many studies highlighting the improved fidelity of *in vitro* air-liquid interface models to their *in vivo* counterparts. Our results regarding the impact of increased oxygen on cell differentiation and function offer a compelling explanation for the efficacy of these air-liquid interface systems and ultimately provide a rationale to allow further improvements to hPSC-based models.

Overall, our findings related to pericellular oxygen concentrations and functional have important implications for cell culture models. First, although we have focussed exclusively on terminally-differentiated cell types, our findings are likely applicable to other cell types. However, it is probable that factors such as differentiation stage and whether the cells are proliferating will affect whether cells in culture experience hypoxia and the cellular response to experimental changes in oxygen tension. Indeed, a key determinant of oxygen consumption at the cell monolayer is cell density or confluence. Our study focuses on terminally-differentiated cell types, which are typically grown to 100% confluence. This is distinct from rapidly proliferative cell-types, such as cancer or immune cells, which proliferate rapidly and are usually grown and studied at pre-confluence. Thus, whether specific cells in culture experience hypoxia and would functionally benefit from increased oxygen needs to be determined on a cell type-to-cell type basis

Second, our study highlights a critical distinction between incubator oxygen levels, and the local oxygen concentration experienced by cells. This suggests that reporting of incubator oxygen % is insufficient, and that measuring pericellular oxygen is required to reveal the actual oxygen concentration experienced by cells. Notably, culturing 3T3-L1 adipocytes in low medium accurately matched *in vivo* oxygen tensions^33, 34^ as well as lactate export^32^. Shifting cells to a more physiological oxygen tension correlated with improved adipocyte function (Figure 3). Since cells *in vivo* exist on a spectrum of oxygen tensions^6^, we suggest that the best reference point for optimal oxygen tension for a specific cell-type is to match the pericellular oxygen to that of the relevant tissue *in vivo*.

Finally, the powerful effects of adjusting oxygen tension on cell phenotypes highlights the need for researchers to control and report on factors that impact oxygen availability (e.g. medium volumes, cell densities) to ensure reproducibility. For instance, acute changes in medium volumes (e.g. adding more medium to cells over the weekend, or reducing medium volumes during assays to save on expensive reagents) may lead to profound changes in experimental findings.

Our findings ultimately highlight that cell culture is a state of variable oxygen tension depending on factors such as oxygen consumption by cells and limitations of oxygen diffusion through the media column. Manipulating oxygen levels can cause dramatic effects on many aspects of cellular metabolism and function. As such, these findings complement recent data on the use of more physiological medium^4, 5^, with potentially important implications for the translatability of both cell and tissue culture models to *in vivo* settings.

## Supporting information

Table S6: scWAT RNAseq Fast Gene Set analysis (KEGG pathways)

Table S1: 3T3-L1 adipocyte 4/16h metabolomics

Table S2: 3T3-L1 adipocyte RNAseq differentially expressed genes

Table S3: 3T3-L1 adipocyte RNAseq VIPER transcription factor analysis

Table S4: 3T3-L1 adipocyte RNAseq Fast Gene Set analysis (KEGG pathways)

Table S5: scWAT RNAseq differentially expressed genes

## ACKNOWLEDGMENTS

These studies were supported by the Wellcome-MRC Institute of Metabolic Science (IMS) Metabolic Research Laboratories, Imaging Core (Wellcome Trust Major Award (208363/Z/17/Z)), and the MRC MDU Mouse Biochemistry Laboratory (MC_UU_00014/5). RNAseq was performed by the IMS Genomics and transcriptomics core facility and supported by the UK MRC Metabolic Disease Unit (MRC_MC_UU_00014/5) and a Wellcome Trust Major Award (208363/Z/17/Z). Schematic illustrations (Figures 1D and S3C) were created with BioRender. S.V. was supported by BHF (RG/18/7/33636). O.J.C. was supported by a Wellcome Trust PhD studentship. I.K. was supported by a Medical Research Council (MRC) PhD studentship. C.P. was supported by a BBSRC project grant (BB/W005905/1). A.J.M was supported by BBSRC (BB/F016581/1) and BHF (FS/17/61/33473). D.C.G. is funded by a Sir Henry Dale Fellowship from the Wellcome Trust/Royal Society (210481). J.E.H. was supported by a Snow Medical Fellowship. The L.V. lab is funded by the ERC advanced grant New-Chol and the core support grant from the Wellcome Trust and Medical Research Council (MRC) of the Wellcome–Medical Research Council Cambridge Stem Cell Institute. For K.H.F.-W. the work was supported in part by DOD-W81XWH-19-1-0213. C.F. and M.Y. were supported by the MRC Core award (MRC_MC_UU_12022/6). A.V-P. was supported by BHF (RG/18/7/33636) and MRC (MC_UU_12012/2). D.J.F. was supported by a Medical Research Council Career Development Award (MR/S007091/1) and a Wellcome Institution Strategic Support Fund award (204845/Z/16/Z).

## AUTHOR CONTRIBUTIONS

Conceptualisation (S.V., D.M.N., and D.J.F.), methodology (J.T., S.V., D.M.N., A.S.S., and D.J.F.), formal analysis (J.T., S.V., M.Y., and I.K.), investigation (J.T., S.V., D.M.N., O.J.C., M.Y., C.G., F.L., I.K., J.R.K., C.P., M.D., H.J.M.B., J.A.H., A.P.S., E.N., M.M., S.C., E.P., A.J.M., D.C.G., J.E.H., L.V., K.H.F-W., and D.J.F.), writing of manuscript (J.T., S.V., and D.J.F., with inputs from all authors), visualisation (J.T., S.V., M.Y., K.H.F-W., and J.E.H.), supervision (S.V., E.P., A.J.M., D.C.G., J.E.H., L.V., C.F., A.V-P., and D.J.F.), project administration (J.T., S.V., and D.J.F.).

## RIGHTS RETENTION STATEMENT

This work was funded by a UKRI grant (MR/S007091/1). For the purpose of open access, the author has applied a Creative Commons Attribution (CC BY) licence to any Author Accepted Manuscript version arising.

## RESOURCE AVAILABILITY

### Lead contact

Further information and requests for resources and reagents should be directed to and will be fulfilled by the lead contact, Daniel Fazakerley.

### Materials availability

This study did not generate new unique reagents.

### Data and code availability

All the fastq files and relevant metadata are available through Array Express (www.ebi.ac.uk/arrayexpress), with accession numbers E-MTAB-12298 and E-MTAB-12299. Metabolomics data are currently being deposited to the EMBL-EBI MetaboLights database (DOI: 10.1093/nar/gkz1019, PMID:31691833) with the identifier MTBLS6677 (https://www.ebi.ac.uk/metabolights/MTBLS6677) and will be made available prior to publication. There are no restrictions on data availability. This paper does not report original code. Any additional information required to reanalyze the data reported in this paper is available from the lead contact upon request.

## METHOD DETAILS

### Cell culture of 3T3-L1 adipocytes

3T3-L1 fibroblasts were cultured in high glucose DMEM supplemented with 10% foetal bovine serum (FBS) and 2 mM glutamax at 37°C in 10% CO_2_. For hypoxia experiments, cells were cultured at 37°C in 5% O_2_, 5% CO_2_. For differentiation into adipocytes, fibroblasts were cultured in 10% FBS-supplemented DMEM containing an adipogenic cocktail (350 nM insulin, 0.5 mM 3-isobutyl-1-methylxanthine, and 250 nM dexamethasone for 3 d, followed by 3 d in DMEM containing 10% FBS and 350 nM insulin. Differentiated adipocytes were maintained in 10% FBS-supplemented DMEM. Adipocytes were used for experiments 9–10 d after the onset of differentiation, with culture medium renewed every 2 days prior to each experiment. For low medium volume treatments, cells were cultured in either 1000 μL (high), 666 μL (mid), 333 μL (low) medium in a 12-well plate, or 500 µL (high), 333 µL (mid), or 167 µL (low) in a 24-well plate (Figure S1A) for the duration specific to each experiment.

### Measurement of extracellular glucose and lactate concentrations

3T3-L1 adipocytes, pBAT, and L6 myotubes were fed with 1 mL (high), 0.67 mL (mid), or 0.33 mL (low) of fresh medium for 16 h. iPSC-derived hepatocytes were fed with 1 mL (high) or 0.5 mL (low) of fresh medium for 24 h. hPSC-derived cardiac organoids were fed with 150 µL (high) or 50 µL (low) of fresh medium for 48 h. For experiments using different starting concentrations of glucose, the appropriate amount of D-(+)-glucose was added to glucose-free DMEM (Sigma #D5030) supplemented with 1 mM sodium pyruvate, 10% FBS, 2 mM glutamax, and 44 mM sodium bicarbonate. Following low medium volume interventions, media was collected from wells and sent for glucose consumption/lactate production analysis at the Core Biochemical Assay Laboratory (Addenbrooke’s Hospital, Cambridge). Naïve medium was used as a baseline. Medium glucose was measured using an adaption of the hexokinase-glucose-6-phosphate dehydrogenase method described by Kunst et al. (1983)^61^ (Siemens Healthcare (product code DF30)). Medium lactate was measured by monitoring absorbance at 340 nm due to NADH production as L-lactate is oxidised to pyruvate by lactate dehydrogenase (Siemens Healthcare (product code DF16)). Calculated glucose use and lactate production were normalised to well areas (except in the case of cardiac organoids). Medium lactate from the cardiac organoids were measured using an in-house enzymatic assay, based on the hydrazine-sink method as described previously^18^. Cardiac data were normalised to account for data variability between batches.

### Measurements of pericellular oxygen concentration and oxygen consumption rate

3T3-L1 adipocytes were cultured in Falcon flat-bottom 96-well microplate (Corning #353072) with 100 µL medium throughout differentiation. Oxygen consumption rates (OCR) and oxygen concentration were continuously measured using Resipher (Lucid Scientific) at 37°C, 10% CO_2_ over 2-3 days starting from 8 d post-differentiation. The Resipher oxygen sensing lid contains micro probes with optical oxygen sensors that scan between 0.55–0.95 mm above the cells to measure the oxygen concentration gradient within the medium column. Baseline OCR and oxygen concentrations under 100 µL medium were measured for 4-6 h in each well. Then, medium volumes were changed and replenished every 24 h. 100 nM antimycin A treatment and trypsinised cells for medium-only wells were used as controls. Data were analysed using the Resipher web application.

### Relative measurements of pericellular oxygen concentration

3T3-L1 cells were differentiated in 96-well OxoPlates (PreSens), which have optical oxygen sensors integrated at the bottom of each well. Prior to starting the experiment, different volumes of fresh medium were added to each well, and the plate immediately placed into the Tecan Spark 10M Plate Reader set to 37°C, 10% CO2 for 24 h. The filters were set up and oxygen levels calculated according to the manufacturer’s protocol. Relative oxygen levels were normalised to the 100 µL condition.

### 3T3-L1 culture in gas-permeable (Lumox) plates

Differentiated 3T3-L1 cells were reseeded on day 6 post-differentiation into 24-well plates (control), or 24-well Lumox plates (Sarstedt), which consist of a black polystyrene frame with a transparent base made from a thin (50 μm), gas-permeable film. The medium volume was kept at 500 μL throughout the differentiation protocol. At day 9 post-differentiation, 500 μL fresh medium was added and collected after 16 h for glucose and lactate quantification, or cells were scraped for lipid extraction as detailed in the other sections.

### Deuterium tracing and lipid extraction

Prior to lipid extraction, cells were cultured in DMEM containing 8% ^2^H (deuterium)-labelled water, in a 37°C, 10% CO_2_ incubator containing 8% ^2^H-labelled water for 16 h. Cells were detached with 100 μL ice-cold PBS and lipids extracted and derivatised as fatty acid methyl esters (FAMEs). Lipids were extracted according to a modified Folch method^62^ using 50 μM tridecanoic-d25 acid (Cambridge Isotope Laboratories, Inc.) as an internal standard. 1 mL of HPLC-grade chloroform/methanol) 2:1 v/v mixture was added to cell samples in a glass vial. Samples were homogenised by vortexing for 5 min. 200 μL H_2_O was added to each sample before vortexing again for 5 min and centrifuging at 4000 x g for 10 min. 600 μL of the lower lipid fraction was transferred to a 7 mL glass tube. A second extraction was performed by 11 adding 600 μL fresh chloroform followed by vortexing and centrifugation as above. Another 600 μL of lower lipid fraction was collected and pooled with the first 600 μL fraction (total = 1200 μL). The collected organic fraction was dried under nitrogen steam and stored at −80°C for subsequent gas chromatography-mass spectrometry (GC-MS) analysis.

### Fatty acid methyl ester derivatisation and GC-MS analysis

Dried lipids were dissolved in a mixture of 410 μL methanol, 375 μL chloroform, and 90 μL 14% BF3 for the FAMEs, and incubated at 80°C for 90 min. 500 μL H2O and 1000 μL hexane were then added to each sample, homogenised by vortexing briefly, and centrifuged at 2000 rpm for 5 min at room temperature. The organic upper layer was then transferred to an autosampler vial for GC-MS analysis.

GC-MS was performed with Agilent 7890B gas chromatography system linked to Agilent 5977A mass spectrometer, using AS3000 autosampler and data was acquired using MassHunter Workstation Software. A TR-FAME column (length: 30 m, inter diameter: 0.25 mm, film size: 0.25 μm, 260M142P, Thermofisher Scientific) was used with helium as carrier gas. Inlet temperature was set at 230°C. Dried FAME samples were re-suspended in 200 μL HPLC-grade n-Hexane. 1 μl of this solution was injected for analysis. The oven programme used for separation was as follows: 100°C hold for 2 min, ramp at 25°C/min to 150°C, ramp at 2.5°C/min to 162°C and hold for 3.8 min, ramp at 4.5°C/min to 173°C and hold for 5 min, ramp at 5°C/min to 210°C, ramp at 40°C/min to 230°C and hold for 0.5 min. Carrier gas flow was set to constant 1.5 mL/min. If the height of any FAME peaks exceeded 108 units, sample was re-injected with 10:1 – 100:1 split ratio. Identification of FAME peaks was based on retention time and made by comparison with those in external standards (Food industry FAME mix, 35077, Restek).

Peak integration and quantification were performed using MassHunter Workstation Quantitative Analysis software (version B.07.00, Agilent). Specific high-abundance ions from total ion chromatograms were chosen to calculate each fatty acid peak.

### Mass isotopomer distribution analysis (MIDA)

To calculate *de novo* lipogenesis rates from fatty acids, MIDA was performed according to the following protocol. In order to perform the analyses we extracted the M+0 to M+4 ions (e.g. for palmitate methyl ester 12 (m/z 270-274)). From these we calculated the fractional concentration of each ion. All equations below use fractional concentrations.

We determined a theoretical distribution for a newly synthesised molecule that was dependent on the precursor labelling pool p, whereby p was the fraction of deuterated water (i.e. 0.08). Firstly, we calculated the isotopomer pattern of each fatty acid caused by the presence of additional 2H atoms in the molecule. The number of available sites in a fatty acid that could be deuterated, N, was determined based on model fit (e.g. for palmitate in 3T3-L1 adipocytes, N = 14). Ions dependent on deuterium incorporation were called M’. For each ion M’0-M’4 the following equation gave the expected relative abundance:

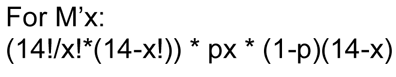

The pattern of labelling based on deuterium was then corrected for the presence of naturally occurring oxygen and carbon isotopes using the following equations for ions M0n-M4n, where Mxn stood for the ion of a newly synthesised fatty acid molecule:

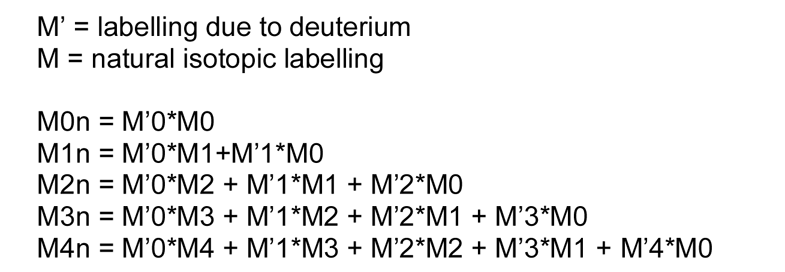

These equations provided us with the ion pattern for a newly synthesised fatty acid M0n-M4n.

To determine the relative contribution of newly synthesised fatty acids and existing fatty acids within the cell we set the following equations, where M0n is newly synthesised, M0 was endogenous pre-existing fatty acids and M0obs was the observed M0 fractional concentration of the ion measured using the mass spectrometer. M0obs’ was the calculated fraction of M0 based on combining newly synthesised and pre-existing fatty acid as follows:

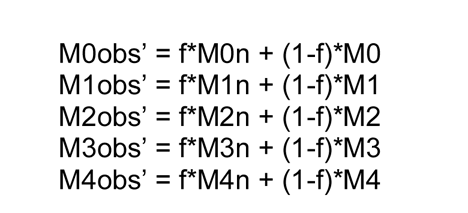

We then calculated M0obs-M0obs’ through to M4obs-M4obs’ and calculated the sum of squares for these 4 equations. We used the GRG Non-Linear Engine of the Solver function of excel to minimise the sum of squares of M0obs-M0obs’+…M4obs-M4obs’ by changing the values of p and f. This enabled us to calculate the fractional synthesis rate for each fatty acid. The N number was adjusted until the p was 0.08 (8% deuterated water) which was experimentally set, thus providing the N number. The amount of newly synthesised palmitate was calculated by multiplying the f value by the concentration of the fatty acid in the well.

### Measurements of oxygen consumption rate in a closed system

High-resolution O_2_ consumption measurements were conducted using the Oroboros Oxygraph-2K (Oroboros Instruments, Innsbruck, Austria) in intact and digitonin-permeabilised cells. For cell experiment, cells were centrifuged at 300 x g for 7 min at room-temperature, washed in PBS, centrifuged once more and then suspended in assay medium at a cell concentration of ∼1 x 10^6^ viable cells/mL. For experiments designed to assess basal respiratory kinetics in intact cells, differentiated 3T3-L1 adipocytes were suspended in bicarbonate-free DMEM, supplemented with Glutamax (glucose 4.5 g/L). Respiration and oxygen tension were continuously evaluated until the cells consumed all the available oxygen. For experiments in permeabilized cells, intact 3T3-L1 adipocytes, exposed to low or high medium for 16 h, were suspended in respiration buffer. Respiration buffer consisted of potassium-MES (105 mM; pH 7.2), KCl (30 mM), KH_2_PO_4_ (10 mM), MgCl_2_ (5 mM), EGTA (1 mM), and BSA (2.5 g/L). Following basal respiration, cells were permeabilised with digitonin (0.02 mg/mL). Physiological ATP free energy (-54.16 kJ/mol) was applied via the creatine kinase (CK) clamp^63^. Cytochrome *c* (0.01 mM) was added to assess the integrity of the outer mitochondrial membrane, followed by sequential additions of carbon substrates and respiratory inhibitors [pyruvate/malate (Pyr/Mal 5mM/1mM); Glutamate (10 mM); octanoyl-carnitine (Oct 0.2 mM); rotenone (Rot 0.5 µM); Succinate (10 mM); Malonate (10 mM); Calcium (0.6 µM); glycerol-3-phosphate (G3P 10 mM); Antimycin (0.5 µM)]. Data were normalised to total protein.

### Immunoblotting

Cells were washed three times in ice-cold PBS and lysed in RIPA buffer containing protease and phosphatase Inhibitors (ThermoFisher). Scraped lysates were then sonicated and centrifuged at 16,000 x g at 4 °C for 30 min. Protein concentration of the supernatant was quantified using the BCA Assay (Thermo Scientific). Lysates were diluted in 4x Laemmli Sample Buffer (Bio-Rad), reduced with TCEP (ThermoFisher) and heated at 37°C for 30 min (Figures 2C and S2A; GLUT1, GLUT4, mitochondrial respiratory complexes), or 65°C for 10 min (Figure 2D; pPDH, PDH). 10-20 μg of protein was resolved by SDS-PAGE and transferred to nitrocellulose membranes (Bio-Rad). Membranes were blocked in 5% skim milk powder in Tris-buffered saline for 1 h at room temperature, followed by an overnight incubation at 4°C with specific primary antibody solutions. Subsequently, membranes were incubated with the appropriate secondary antibodies for 1 h at room temperature before signals were detected using enhanced chemiluminescence (ECL) (Thermo Scientific) on the Chemidoc MP (Bio-Rad). Bands were quantified using ImageJ2. pPDH (1:2000, #31866 Cell Signaling Technology), PDH (1:2000, #2784 Cell Signaling Technology), GLUT4 (1:1000, a gift from F. Koumanov, University of Bath and affinity-purified in-house), GLUT1 (1:1000, #12939 Cell Signaling Technology), α-tubulin (1:1000, #T9026 Sigma Aldrich), OxPhos Rodent WB Antibody Cocktail (1:250, #45-8099 Invitrogen) and β-actin (1:1000, #8457 Cell Signaling Technology) antibodies were used.

### Immunoblotting

Cells were washed three times in ice-cold PBS and lysed in 2% SDS containing protease and phosphatase Inhibitors (ThermoFisher), 500 µM CoCl_2_, and 10 µM MG132. Scraped lysates were then sonicated and centrifuged at 16,000 x g at room temperature for 30 min. Protein concentration of the supernatant was quantified using the BCA Assay (Thermo Scientific). Lysates were diluted in 4x Laemmli Sample Buffer (Bio-Rad), reduced with TCEP (ThermoFisher) and heated at 95°C for 10 min. 40 μg of protein was resolved by SDS-PAGE and transferred to nitrocellulose membranes (Bio-Rad). Membranes were blocked in 5% skim milk powder in Tris-buffered saline for 1 h at room temperature, followed by an overnight incubation at 4°C with anti-HIF1α (1:500, #14179 Cell Signaling Technology). Subsequently, membranes were incubated with the goat anti-rabbit IgG for 1 h at room temperature before signals were detected using ECL (Thermo Scientific) on the Chemidoc MP (Bio-Rad). Bands were quantified using ImageJ2.

### Assessment of PDH flux

PDH flux was assessed using the FASA package implemented in Matlab^64^. FASA was performed on isotope corrected isotopomer distributions for palmitate (M0-M16). Isotope correction was performed using the IsoCorR package implemented in R. The value “D” from the FASA analysis was the proportion of lipogenic acetyl-CoA derived from glucose that was used to produce palmitate.

### ^13^C-glucose tracing and metabolite extraction

Prior to the experiment, 3T3-L1 adipocytes were washed once with glucose-free DMEM and labelled with 25 mM 13C-glucose in DMEM for either 4 h or 16 h. Then, medium was removed from the wells and cells were washed twice with room temperature PBS and placed on dry ice. 250 μL metabolite extraction buffer (50% methanol, 30% acetonitrile, 20% ultrapure water, 5 µM valine-d8) was added to each well of a 12-well plate and incubated for 5 min on a dry ice-methanol bath to lyse the cell membranes. After that, extracts were scraped and mixed at 4°C for 15 min in a thermomixer at 2000 rpm. After final centrifugation at max speed for 20 min at 4°C, 80 μL of the supernatant from each sample was transferred into labelled LC-MS vials.

### LC-MS analysis

Hydrophilic separation of metabolites was achieved using a Millipore Sequant ZIC-pHILIC analytical column (5 µm, 2.1 × 150 mm) equipped with a 2.1 × 20 mm guard column (both 5 mm particle size) with a binary solvent system. Solvent A was 20 mM ammonium carbonate, 0.05% ammonium hydroxide; Solvent B was acetonitrile. The column oven and autosampler tray were held at 40 °C and 4 °C, respectively. The chromatographic gradient was run at a flow rate of 0.200 mL/min as follows: 0–2 min: 80% B; 2-17 min: linear gradient from 80% B to 20% B; 17-17.1 min: linear gradient from 20% B to 80% B; 17.1-23 min: hold at 80% B. Samples were randomised and the injection volume was 5 µl. A pooled quality control (QC) sample was generated from an equal mixture of all individual samples and analysed interspersed at regular intervals.

Metabolites were measured with Vanquish Horizon UHPLC coupled to an Orbitrap Exploris 240 mass spectrometer (both Thermo Fisher Scientific) via a heated electrospray ionisation source. The spray voltages were set to +3.5kV/-2.8 kV, RF lens value at 70, the heated capillary held at 320 °C, and the auxiliary gas heater held at 280 °C. The flow rate for sheath gas, aux gas and sweep gas were set to 40, 15 and 0, respectively. For MS1 scans, mass range was set to *m/z*=70-900, AGC target set to standard and maximum injection time (IT) set to auto. Data acquisition for experimental samples used full scan mode with polarity switching at an Orbitrap resolution of 120000. Data acquisition for untargeted metabolite identification was performed using the AcquireX Deep Scan workflow, an iterative data-dependent acquisition (DDA) strategy using multiple injections of the pooled sample. DDA full scan-ddMS2 method for AcquireX workflow used the following parameters: full scan resolution was set to 60000, fragmentation resolution to 30000, fragmentation intensity threshold to 5.0e3. Dynamic exclusion was enabled after 1 time and exclusion duration was 10s. Mass tolerance was set to 5ppm. Isolation window was set to 1.2 *m/z*. Normalised HCD collision energies were set to stepped mode with values at 30, 50, 150. Fragmentation scan range was set to auto, AGC target at standard and max IT at auto. Mild trapping was enabled.

Metabolite identification was performed in the Compound Discoverer software (v 3.2, Thermo Fisher Scientific). Metabolite identities were confirmed using the following parameters: (1) precursor ion m/z was matched within 5 ppm of theoretical mass predicted by the chemical formula; (2) fragment ions were matched within 5 pm to an in-house spectral library of authentic compound standards analysed with the same ddMS2 method with a best match score of over 70; (3) the retention time of metabolites was within 5% of the retention time of a purified standard run with the same chromatographic method. Chromatogram review and peak area integration were performed using the Tracefinder software (v 5.0, Thermo Fisher Scientific) and the peak area for each detected metabolite was normalised against the total ion count (TIC) of that sample to correct any variations introduced from sample handling to instrument analysis. The normalised areas were used as variables for further statistical data analysis.

For ^13^C_6_-glucose tracing analysis, the theoretical masses of ^13^C-labelled isotopes were calculated and added to a library of predicted isotopes in Tracefinder 5.0. These masses were then searched with a 5-ppm tolerance and integrated only if the peak apex showed less than 1% deviation in retention time from the [U-^12^C] monoisotopic mass in the same chromatogram. The raw data obtained for each isotopologue were then corrected for natural isotope abundances using the AccuCor algorithm (https://github.com/lparsons/accucor) before further statistical analysis.

### ^3^H-2-deoxyglucose uptake measurements

3T3-L1 adipocytes were differentiated in 24-well plates with 500 µL medium throughout differentiation. 48 h prior to the assay, medium volumes were changed to either 500 µL (high) or 167 µL (low). On the day of the assay, cells were serum-starved for 2 h with either 500 or 167 µL DMEM containing 0.2% BSA at 37°C, 10% CO_2_. Following 2 h serum-starvation, cells were washed and incubated in pre-warmed Krebs–Ringer phosphate (KRP) buffer containing 0.2% bovine serum albumin (BSA, Bovostar, Bovogen) (KRP buffer; 0.6 mM Na_2_HPO_4_, 0.4 mM NaH_2_PO_4_, 120 mM NaCl, 6 mM KCl, 1 mM CaCl_2_, 1.2 mM MgSO_4_ and 12.5 mM Hepes (pH 7.4)) for 10 min. Again, KRP volumes were maintained at either 500 or 167 µL, and 200 µM indinavir^65^ was added to at this point where indicated. Adipocytes were then stimulated with 100 nM insulin for 20 min. To determine non-specific glucose uptake, 25 μM cytochalasin B (in ethanol, Sigma Aldrich) was added to control wells before addition of 2-[^3^H]deoxyglucose (2-DG) (PerkinElmer). During the final 5 min 2-DG (0.25 μCi, 50 μM) was added to cells to measure steady-state rates of 2-DG uptake. Note that the volumes of 2-DG added were altered to account for different medium volumes. Cells were then moved to ice,washed with ice-cold PBS, and solubilised in PBS containing 1% (v/v) Triton X-100. Tracer uptake was quantified by liquid scintillation counting on the TriCarb 2900TR (PerkinElmer) and data normalised for protein content.

### qRT-PCR

3T3-L1 adipocytes were cultured using 1 mL medium (12-well plate) throughout differentiation. Medium volumes were changed to either 1 mL or 0.33 mL 16 h prior to the assay. Hepatocytes were cultured in either 1 mL or 0.5 mL medium (12-well plate) throughout differentiation, and changed to either 1 mL or 0.5 mL 24 h prior to the assay. RNA extractions were performed using the RNeasy Mini kit (Qiagen 1152 #74104). Concentrations of RNA samples were quantified using NanoDrop. cDNA synthesis from 500 ng RNA was performed using the GoScript Reverse Transcriptase kit (Promega #A2801). Real-time (RT)-polymerase chain reaction (PCR) was performed with SYBR Green Master Mix on the ABI QuantStudio 5.

### Subcutaneous white adipose tissue (scWAT) from normoxic and hypoxic mice

Animal work was carried out in accordance with United Kingdom Home Office regulations under the Animals in Scientific Procedures (1986) Act and underwent review by the University of Cambridge Animal Welfare and Ethical Review Board. All procedures involving live animals were carried out by a personal licence holder in accordance with these regulations.

Subcutaneous adipose tissue (scWAT) was collected from 80-day old male mice exposed to either 10% O_2_ or maintained under normoxic atmospheric conditions, from a larger experimental cohort as previously published^66, 67^. 129Ev/Sv mice were housed in conventional cages from birth in a temperature-(23°C) and humidity-controlled environment with a 12 h photoperiod and given *ad libitum* access to standard rodent chow (RM1(E), Special Diet Services, UK) and water. At 7 weeks of age, mice were randomly assigned to remain under normoxic conditions, or transferred to a normobaric hypoxia chamber (10% O_2_; PFI systems, Milton Keynes, UK). Mice were maintained in hypoxia or normoxia for 28 days, after which they were killed by cervical dislocation. The scWAT was removed from the inguinal region, snap frozen and stored at −80°C until further analysis. The inguinal lymph node was removed from scWAT depots prior to RNA-seq analysis.

### RNA extraction and sequencing

3T3-L1 adipocytes were seeded and differentiated in 12-well plates. RNA extraction from adipocytes was performed using the RNeasy Mini Kit (Qiagen 1152 #74104).

RNA extraction from mice scWAT was performed by homogenisation in RNA STAT-60. Samples were broken up using a pestle and mortar under liquid nitrogen. The broken pieces were then put in 2 mL screw-top tubes with Lysing Matrix D (MP Biomedicals). 1 mL STAT-60 was added to the tissue samples, followed by homogenisation using Precellys 24 Touch. The homogenate was then transferred to 1.5 mL eppendorfs. Tubes were spun at 13,000 rpm for 5 min at 4°C to remove debris. The supernatant was removed and transferred to fresh tubes containing 200 µL of Chloroform. Samples were vortexed at maximum speed for 15 s, and spun at 12,000 g for 15 min at 4°C. A clear aqueous supernatant of around 600 µL for every 1 mL of STAT-60 added will have formed. The aqueous phase was transferred to fresh tubes containing 500 µL of isopropanol. Tubes were mixed by inversion several times and left at −20°C overnight. After overnight incubation, the precipitates were spun at 12,000 g for 10 min at 4°C to pellet RNA. After removing the supernatant, 1 mL of cold 70% ethanol (stored at − 20°C) was added to the pellet and washed by vortexing briefly. Samples were then spun at 8000 g for 5 min at 4°C before discarding the supernatant. Pellets were air-dried for 10-15 min on ice until transparent. Nuclease-free water was added to the pellet and incubated at 60°C for 10 min. Samples were placed on ice and re-suspended by pipetting up and down.

RNA samples from both 3T3-L1 adipocytes and mice scWAT were quantified using nanodrop. 1 μg of total RNA was quality checked (RIN >7) using an Agilent Bioanalyser 2100 system and then used to construct barcoded sequencing libraries with Illumina’s TruSeq Stranded mRNA Library Prep Kit following manufacturer’s instruction. All the libraries were then multiplexed and sequenced on one lane of Illumina NovaSeq6000 at PE50 at the CRUK Cambridge Institute Genomics Core Facility.

### RNAseq data analysis

The NGS data were processed through a customised pipeline. FastQC software (v. 0.11.9) was used for generating quality-control reports of individual FASTQ files (http://www.bioinformatics.babraham.ac.uk/projects/fastqc). RNA sequencing reads were aligned to the *mus musculus* (mouse) GRCm38 genome using hisat2 (v2.1.0)^68^. HTseq-count (v 0.11.1)^69^ was then used for gene counting and DESeq2 (v3.15)^70^ for differential gene expression analysis (Wald Test). The raw p values were adjusted by the Benjamini-Hochberg procedure to control the False Discovery Rate (FDR). Pathway enrichment analysis was performed with FGSEA (fast gene set enrichment analysis) using pre-ranked gene lists and pathways from the KEGG database^71, 72^.

For the inference of the upstream transcriptional regulators we used VIPER (Virtual Inference of Protein-activity by Enriched Regulon analysis)^73^. The estimation of their activation/inhibition status is based on the differential expression of their known target genes and is represented by a positive or negative NES (normalised enrichment score), respectively. The network that links the transcriptional regulators with their target genes was derived from DoRothEA (https://saezlab.github.io/dorothea/)^74^.

### Measurement of extracellular leptin and adiponectin

3T3-L1 adipocytes were cultured as described above. Cells were transferred to low or high medium at the volumes stated above for 48 h, before media was collected and analysed by the Core Biochemical Assay Laboratory (Addenbrooke’s Hospital, Cambridge). Leptin concentration was determined using the MesoScale Discovery Mouse Leptin Kit (K152BYC-2, Rockville, MD, USA), and adiponectin concentration using the MesoScale Discovery Mouse Adiponectin Kit (K152BXC, Rockville, MD, USA) according to manufacturer’s instructions.

### Immunofluorescence analysis of GLUT4 translocation in 3T3-L1 adipocytes

Cells were differentiated in black CellCarrier-96 Ultra Microplates (PerkinElmer). Cells were cultured in specified medium volumes for 48 h prior to assay and medium volumes were kept the same throughout the assay until cells were fixed. Cells were serum-starved in DMEM containing 0.2% BSA for 2 h at 37°C, 10% CO_2_, and then stimulated with 0.5 nM or 100 nM insulin for 20 min. Cells were then washed in an ice-cold PBS bath and instantly fixed with 4% paraformaldehyde on ice for 5 min, followed by 15 min at room temperature. Cells were then quenched with 50 mM glycine for 10 min, then washed twice with PBS at room temperature, and incubated in blocking buffer (5% normal swine serum in PBS) for 20 min. After blocking, cells were incubated with anti-GLUT4 (LM048) (2 µg/mL, Integral Molecular, PA, USA) and Lectin-FITC conjugate (Sigma #L4895) for 1 h at room temperature. This antibody recognises an external epitope on GLUT4, and so staining in non-permeabilised cells will only label GLUT4 present in the plasma membrane^75^. Then, cells were washed in PBS and incubated with anti-human Alexa-647 (1:500, ThermoFisher), and Hoechst 33342 (1:5000) room temperature. After 1 h, the plates were washed three times in PBS, then stored and imaged in PBS, 5% glycerol and 2.5% Dabco. Imaging was performed with the Opera Phenix (Perkin Elmer) using a 20x NA1.0 water immersion objective. Nine fields of view were imaged per well and analysed using a custom pipeline in Harmony High-Content Imaging and Analysis Software (Perkin Elmer). Cell plasma membrane regions were defined using the lectin-FITC signal and the anti-GLUT4 signal within this region calculated as a measure of GLUT4 translocation.

### CL dose response and lipolysis assay

3T3-L1 adipocytes were cultured as described above. All lipolysis experiments were performed on day 10 following differentiation. Medium volume interventions were carried out 48 h prior to the experiment, and cells were kept in the appropriate medium volumes (high or low) for the duration of the lipolysis experiments. Cells were serum-starved in DMEM containing 0.2% BSA for 2 h at 37°C, 10% CO_2_, and then stimulated with the appropriate agonist (insulin or CL316,243 (Tocris)) in KRP buffer for 30 min at 37°C. Glycerol release into the KRP buffer was determined using the free glycerol reagent (Sigma, #F6428) and absorbance measured at 595 nm. Glycerol release was normalised to cellular protein content (determined by BCA) of cells lysed in PBS containing 1% triton.

### Cell culture of iPSC-derived hepatocytes

Human induced pluripotent stem cells (iPSCs) were maintained on vitronectin XFTM (10 μg/mL, StemCell Technologies)-coated plates and in Essential 8 (E8) medium consisting of DMEM/F12 (Gibco), L-ascorbic acid 2-phosphate (1%), insulin-transferrin-selenium solution (2%, Life Technologies), sodium bicarbonate (0.7%), and Penicillin/Streptomycin (P/S) (1%), freshly supplemented with TGFβ (10 ng/mL, R&D) and FGF2 (12 ng/mL, Qkine). Cells were split every 5-7 days by incubation with 0.5 μM EDTA (Thermo Fisher Scientific) for 4 min at room temperature and clumps were dissociated into small clumps by pipetting. For iPSC differentiation, iPSCs were split into single cells using Accutase (StemCell Technologies) and seeded at a density of 50,000 cells per cm^2^ in E8 media with 10 µM ROCK Inhibitor Y-27632 (Selleckchem). Cells were differentiated towards hepatocytes as previously described^76^ with minor modifications. For foregut specification, cells were incubated for 5 days in RPMI media with 2% B27 and 50 ng/mL Activin (R&D). Cells were differentiated into hepatocytes using HepatoZYME media (Thermo) with 2 mM L-glutamine, 1% P/S, 2% non-essential amino acids (Thermo), 2% chemically defined lipids (Thermo), 30 µg/mL transferrin (Roche), 14 µg/mL insulin (Roche), 20 ng/mL Oncostatin M (R&D) and 50 ng/mL hepatocyte growth factor (R&D). Cells were analysed at day 32 of differentiation. For media volume experiments, cells were differentiated towards hepatocytes in either 0.5 mL or 1 mL of complete HepatoZYME media, and volumes were changed or unchanged for the last 24 h of the experiment.

### CYP3A4 activity assay

CYP3A4 enzymatic activity was measured using the P450 Glo kit (Promega). Cells were incubated with 1:1000 luciferin-IPA in Hepatozyme complete for 1 h at 37°C. 50 µL of cell culture supernatant was mixed with 50 µL detection reagent and incubated 20 min at room temperature in Greiner white 96-well microplates (Sigma-Aldrich). Luminescence was measured in triplicate on a GloMax plate reader. Hepatozyme complete medium was used as background control.

### Immunofluorescence analysis of albumin in iPSC-derived hepatocytes

Cells were fixed in 4% PFA for 15 min at room temperature, washed 3x in PBS and blocked for 1 h in PBS with 10% donkey serum and 0.1% Triton X-100 for permeabilisation. Anti-albumin antibody (Bethyl Labs #A80-229A) was applied in blocking solution overnight at 4°C. Cells were then washed 3x in PBS and incubated in Alexa Fluor 488-conjugated secondary antibody (Life Technologies) in blocking solution, with 3 µM DAPI for 1 h at room temperature. Cells were then washed 3x in PBS and visualised in the plate using a Zeiss inverted 710 confocal microscope.

### hPSC-derived cardiac organoids

Ethical approval for the use of human embryonic stem cells (hPSCs) was obtained from QIMR Berghofer’s Ethics Committee (P2385), and was carried out in accordance with the National Health and Medical Research Council of Australia (NHMRC) regulations. Female HES3 (WiCell) were maintained in mTeSR PLUS (Stem Cell Technologies)/ Matrigel (Millipore) and passaged using T ReLeSR (Stem Cell Technologies). Quality control was performed with karyotyping (G-banding) and mycoplasma testing. hPSC-derived cardiac organoids were generated as recently described^77^.

Cardiomyocyte and stromal cell differentiation was achieved using previously described protocols^78–81^. hPSCs were seeded on Matrigel-coated flasks at 2 x10^4^cells/cm^2^ and cultured in mTeSR-1 for 4 days. To induce cardiac mesoderm, hPSCs were cultured in RPMI B27-medium (RPMI 1640 GlutaMAX+ 2% B27 supplement without insulin, 200μM L-ascorbic acid 2-phosphate sesquimagnesium salt hydrate (Sigma) and 1% penicillin/streptomycin (ThermoFisher Scientific), supplemented with 5 ng/ml BMP-4 (RnD Systems), 9 ng/ml Activin A (RnD Systems), 5 ng/ml FGF-2 (RnD Systems) and 1 μM CHIR99021 (Stemgent or Stem Cell Technologies). Mesoderm induction required daily medium exchanges for 3 days. This was followed by cardiac specification using RPMI B27-containing 5 μM IWP-4 (Stem Cell Technologies) for another 3 days, and then further 7 days using 5 μM IWP-4 RPMI B27+ (RPMI1640 Glutamax + 2% B27 supplement with insulin, 200μM L-ascorbic acid 2-phosphate sesquimagnesium salt hydrate and 1% penicillin/streptomycin) with medium change every 2-3 days. For the final 2 days of differentiation, hPSCs were cultured in RPMI B27+. Harvest of differentiated cardiac cells involved enzymatic digestion, firstly in 0.2% collagenase type I (Sigma) in 20% fetal bovine serum (FBS, ThermoFisher Scientific) in PBS (with Ca^2+^ and Mg^2+^) at 37°C for 1 hour, and secondly in 0.25% trypsin-EDTA at 37°C for 10 minutes. The trypsin was neutralized, and then cells were filtered through a 100 μm mesh cell strainer (BD Biosciences), centrifuged at 300 x g for 3 minutes, and resuspended in α-MEM Glutamax, 10% FBS, 200 μM L-ascorbic acid 2-phosphate sesquimagnesium salt hydrate and 1% penicillin/streptomycin (MEM++++). Previous flow cytometry analysis indicated that differentiated cardiac cells were ∼70% α-actinin^+^/CTNT^+^ cardiomyocytes, ∼30% CD90 stromal cells^80^.

hCO culture inserts were fabricated using SU-8 photolithography and PDMS moulding^78^. Acid-solubilized bovine collagen 1 (Devro) was salt balanced using 10x DMEM (ThermoFisher Scientific) and pH neutralised using 0.1 M NaOH before combining with Matrigel and then the cell suspension on ice. Each hCO contained 5 × 10^4^ cells, a final concentration of 2.6 mg/ml collagen I and 9% Matrigel. 3.5 μL of suspension was pipetted into the hCO culture insert and incubated at 37°C with 5% CO2 for 45 min in order to gel. After gelling, α-MEM GlutaMAX (ThermoFisher Scientific), 10% foetal bovine serum (FBS), 200 μM L-ascorbic acid 2-phosphate sesquimagnesium salt hydrate (Sigma) and 1% Penicillin/Streptomycin (ThermoFisher Scientific) was added. hCO were subsequently cultured in maturation medium (MM)^78^ with medium changes every 2 to 3 days for 5 days (7 days old hCO). To better approximate adult metabolic provisions a ‘weaning medium’ (WM) was utilised. hCO were cultured in WM containing 4% B27 – insulin, 5.5 mM glucose, 1 nM insulin, 200 μM L-ascorbic acid 2-phosphate sesquimagnesium salt hydrate, 1% P/S, 1% GlutaMAX (100x), 33 μg/mL aprotinin and 10 μM palmitate (conjugated to bovine serum albumin in B27) in DMEM without glucose, glutamine and phenol red (ThermoFisher Scientific) with medium changes every 2-3 days.

The elasticity of the Heart Dyno poles enables the contractile properties to be determined via tracking pole deflection, which directly correlates with force^78^. Videos of 10 s were made of each hCO with the Nikon ANDOR WD Revolution Spinning Disk microscope (magnification 4x). While imaging, hCO were incubated at 37°C, 5% CO2 to prevent changes in contractile behaviour. This facilitated the analysis of the contractile properties of the organoids and the production of time-force graphs^78^. Moreover, data was obtained regarding additional important functional parameters including the contraction rate and the activation and relaxation time of the organoids.

## QUANTIFICATION AND STATISTICAL ANALYSIS

All statistical analyses, unless otherwise stated in figure legends, were carried out using GraphPad Prism 9. Two-tailed paired/unpaired Student’s t-tests were used to compare the means between two groups. One/two-way ANOVA with Šidák correction for multiple comparisons was performed for multigroup (at least three) comparisons. Data are presented as mean ± SEM, of at least three independent biological replicates.

## SUPPLEMENTAL TABLES

Table S1: 3T3-L1 adipocyte 4/16h metabolomics

Table S2: 3T3-L1 adipocyte RNAseq differentially expressed genes

Table S3: 3T3-L1 adipocyte RNAseq VIPER transcription factor analysis

Table S4: 3T3-L1 adipocyte RNAseq Fast Gene Set analysis (KEGG pathways)

Table S5: scWAT RNAseq differentially expressed genes

Table S6: scWAT RNAseq Fast Gene Set analysis (KEGG pathways)

