## Supplemental information for "Limited oxygen availability in standard cell culture alters metabolism and function in terminally-differentiated cells"

**A**

| Plate | 12-well | | | 24-well | | | 96-well | | |
|---|---|---|---|---|---|---|---|---|---|
| Well area ($cm^2$) | 3.8 | | | 1.9 | | | 0.32 | | |
| | Medium volume (µL) | Top men. (mm) | Bottom men. (mm) | Medium volume (µL) | Top men. (mm) | Bottom men. (mm) | Medium volume (µL) | Top men. (mm) | Bottom men. (mm) |
| High (standard) | 1000 | 4.7 | 2.4 | 500 | 3.9 | 2.2 | 100 | 3.3 | 2.5 |
| Mid | 666 | 3.8 | 1.7 | 333 | 3.5 | 1.6 | 50 | 2.2 | 1.5 |
| Low | 333 | 3.0 | 0.8 | 167 | 1.9 | 0.8 | 33 | 2.0 | 0.7 |

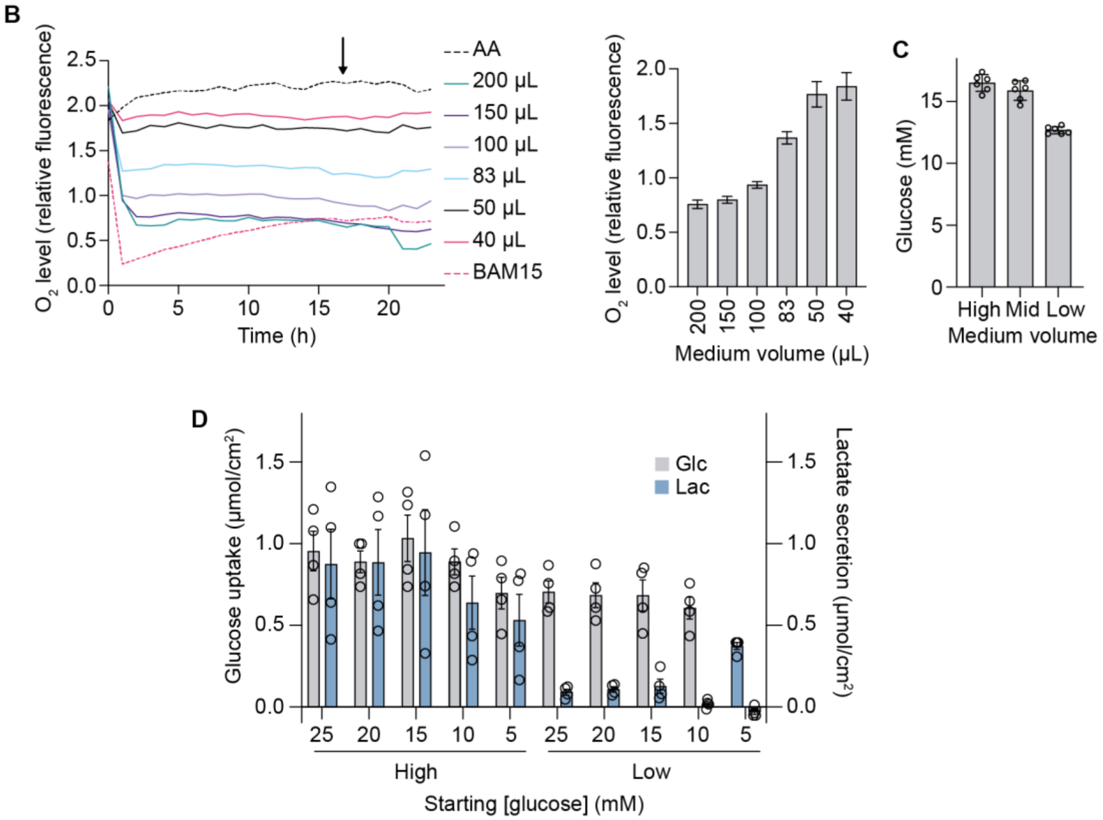


**Fig. S1.**

**Relative oxygen levels at the cell monolayer under different medium volumes**

(**A**) Table of medium volumes used in this study and the corresponding medium heights of both top and bottom menisci. Images of each plate-type containing different medium volumes were used to measure menisci heights. Known well diameters were used to convert menisci heights from pixels to mm. 'High' refers to the standard culture volumes used.

(**B**) Representative trace of fluorescence intensity indicative of pericellular oxygen concentrations under different medium volumes, measured for 24 h in 96-well plates. Bar graph shows relative oxygen levels taken at 16 h (arrow in representative trace) (n = 4 biological replicates). AA, antimycin A.

(**C**) Medium glucose concentration after 16 h of medium volume change in 12-well plates (n = 6 biological replicates).
(**D**) Extracellular medium glucose and lactate measurements after 16 h medium volume change with different starting glucose concentrations in 12-well plates. (n = 4 biological replicates).

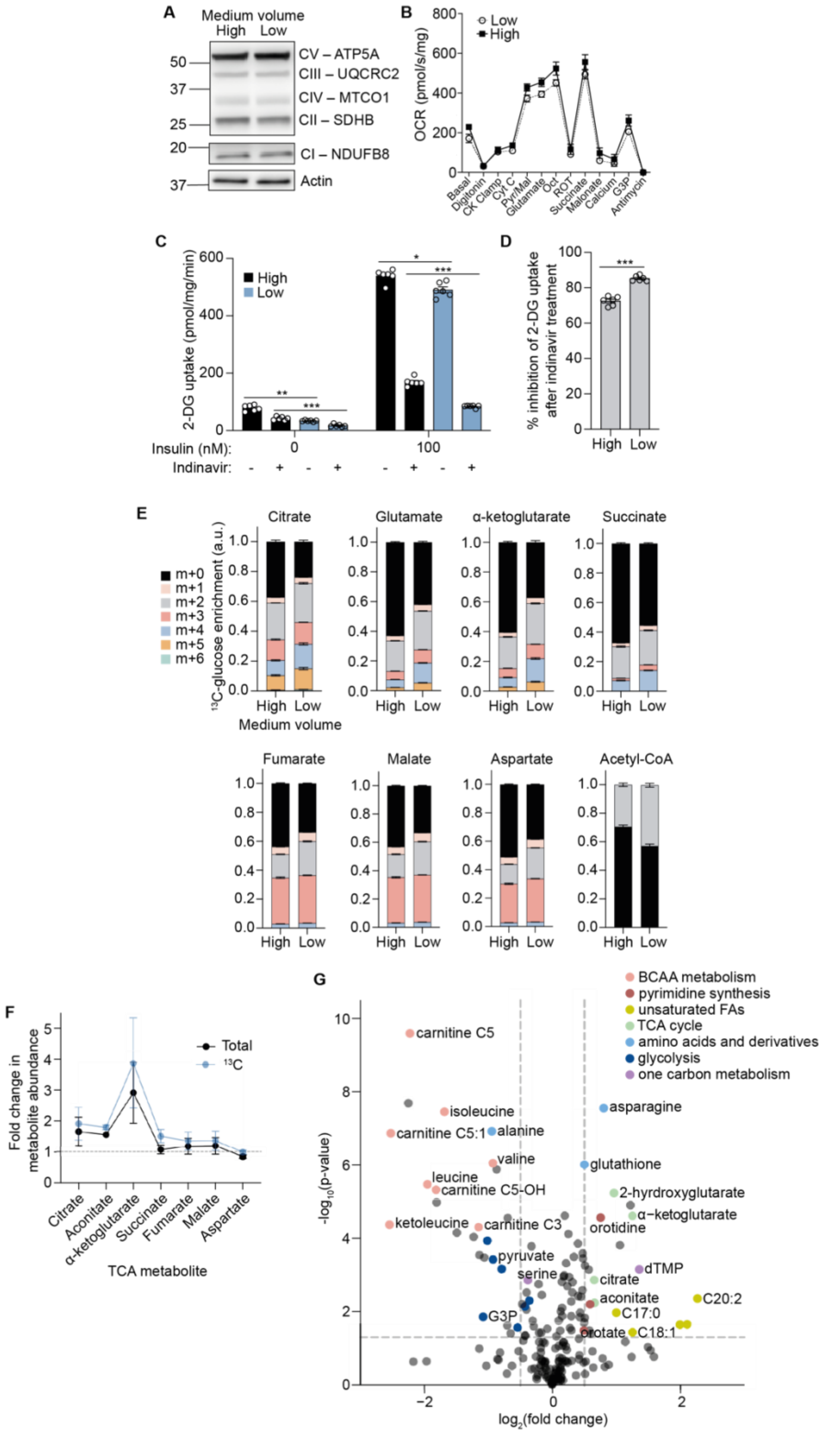

A
Medium volume
High Low
CV – ATP5A
CIII – UQCRC2
CIV – MTCO1
CII – SDHB
CI – NDUFB8
Actin
B
Low
High
OCR (pmol/s/mg)
Basal
Digitonin
CK Clamp
Cyt C
Pyr/Mal
Glutamate
Oct
ROT
Succinate
Malonate
Calcium
G3P
Antimycin
C
2-DG uptake (pmol/mg/min)
High
Low
Insulin (nM):
Indinavir:
D
% inhibition of 2-DG uptake after indinavir treatment
E
Citrate
Glutamate
α-ketoglutarate
Succinate
Fumarate
Malate
Aspartate
Acetyl-CoA
$^{13}$C-glucose enrichment (a.u.)
m+0
m+1
m+2
m+3
m+4
m+5
m+6
Medium volume
F
Fold change in metabolite abundance
Total
$^{13}$C
TCA metabolite
Aconitate
G
BCAA metabolism
pyrimidine synthesis
unsaturated FAs
TCA cycle
amino acids and derivatives
glycolysis
one carbon metabolism
-log$_{10}$(p-value)
log$_2$(fold change)
carnitine C5
isoleucine
carnitine C5:1
alanine
valine
leucine
carnitine C5-OH
ketoleucine
carnitine C3
pyruvate
serine
G3P
asparagine
glutathione
2-hyrdroxyglutarate
α-ketoglutarate
orotidine
dTMP
citrate
aconitate
C20:2
C17:0
orotate
C18:1

**Fig. S2.**

**Increased glucose oxidation is not due to more mitochondrial content or higher glucose uptake in low medium conditions**

(**A**) Western blot of mitochondrial respiratory complexes I–V after 16 h of medium volume change in 12-well plates (n = 3 biological replicates).

(**B**) Oxygen consumption rate (OCR) of permeabilised 3T3-L1 adipocytes upon different substrate stimulation after 16 h of medium volume change (n = 5 biological replicates).

(**C**) 2-deoxyglucose (DG) uptake after insulin stimulation and 200 µM indinavir (GLUT4 inhibitor) treatment. Cells were cultured in high or low medium for 48 h in 24-well plates prior to the experiment (n = 6 biological replicates).

(**D**) Percentage inhibition of 2-DG uptake after indinavir treatment, calculated from the difference between +/- indinavir treated conditions, as a percentage of -indinavir 2-DG uptake upon 100 nM insulin stimulation. Graph shows the percentage of 2-DG uptake that is GLUT4-dependent (i.e. inhibited by indinavir) (n = 6 biological replicates).

(**E**) Fractional abundance of each isotopologue after 4 h medium volume change (n = 6 biological replicates).

(**F**) Fold change of total and $U^{13}C$-glucose labelled TCA metabolite abundance after 16 h medium volume change in 12-well plates (n = 4 biological replicates).

(**G**) Volcano plot of differentially regulated metabolites after 16 h medium volume change. Metabolites of interest which are significantly changed ($p < 0.05$) are highlighted according to their metabolic pathways (n = 6 biological replicates). BCAA, branched-chain amino acid; FA, fatty acid; G3P, glyceraldehyde−3−phosphate.

Data are represented as mean ± SEM. *$p < 0.05$, **$p < 0.01$, ***$p < 0.001$, ****$p < 0.0001$ by two-way ANOVA with Šidák correction for multiple comparisons (C), or by paired two-tailed Student’s t-test (D).

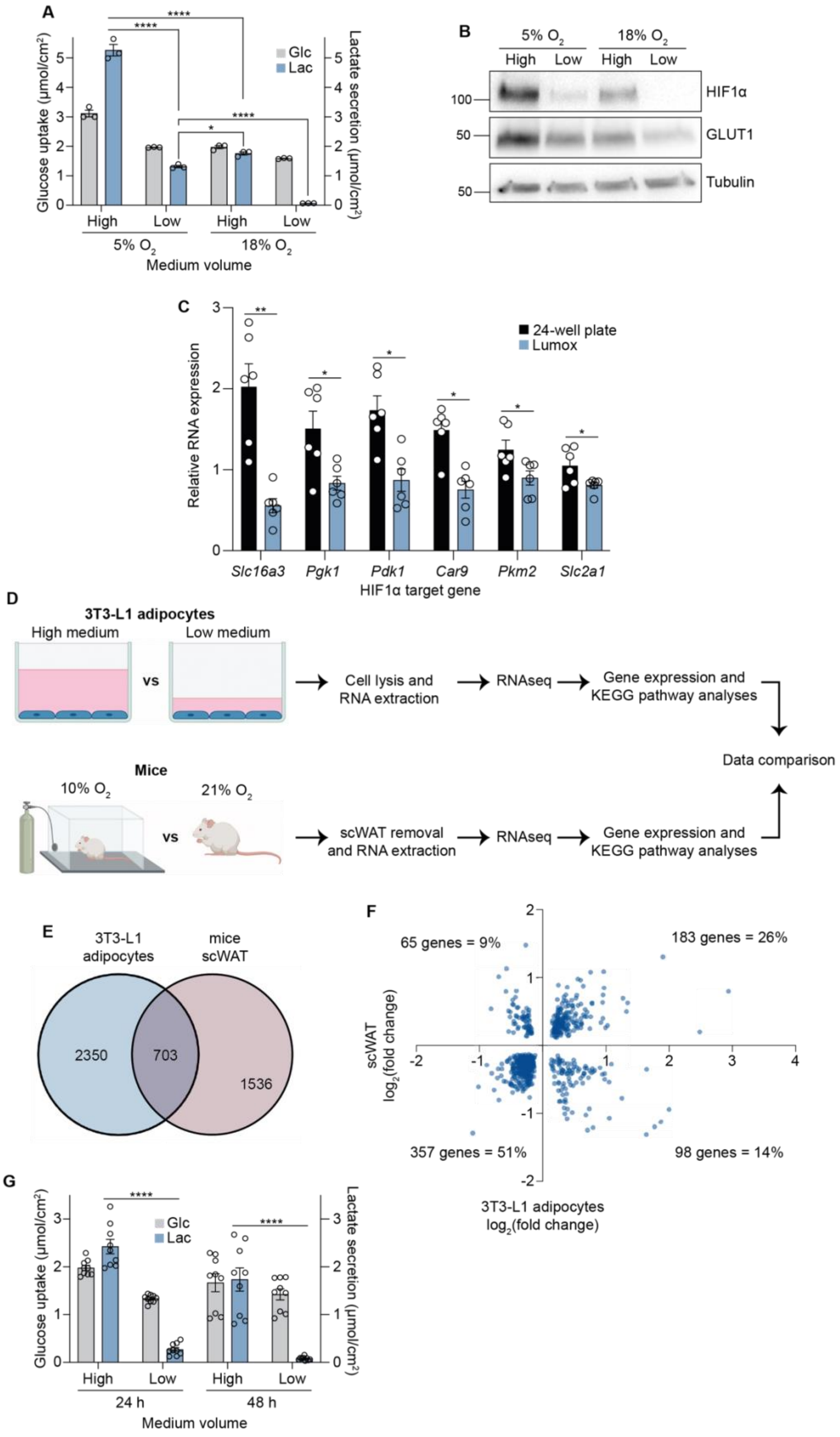
A
Glucose uptake (μmol/cm²)
Lactate secretion (μmol/cm²)
Glc
Lac
****
****
****
*
High
Low
High
Low
5% O2
18% O2
Medium volume
B
5% O2
18% O2
High
Low
High
Low
100
50
50
HIF1α
GLUT1
Tubulin
C
Relative RNA expression
24-well plate
Lumox
**
*
*
*
*
*
Slc16a3
Pgk1
Pdk1
Car9
Pkm2
Slc2a1
HIF1α target gene
D
3T3-L1 adipocytes
High medium
Low medium
vs
Cell lysis and RNA extraction
RNAseq
Gene expression and KEGG pathway analyses
Data comparison
Mice
10% O2
21% O2
vs
scWAT removal and RNA extraction
RNAseq
Gene expression and KEGG pathway analyses
E
3T3-L1 adipocytes
mice scWAT
2350
703
1536
F
65 genes = 9%
183 genes = 26%
357 genes = 51%
98 genes = 14%
scWAT log2(fold change)
3T3-L1 adipocytes log2(fold change)
G
Glucose uptake (μmol/cm²)
Lactate secretion (μmol/cm²)
Glc
Lac
****
****
High
Low
High
Low
24 h
48 h
Medium volume

**Fig. S3.**

**Increasing oxygen availability reduces HIF1α transcriptional activity**

(**A**) Extracellular medium glucose and lactate measurements after 16 h medium volume change (12-well plate) in 5% or 18% oxygen incubators. (n = 3 biological replicates).

(**B**) Western blot of hypoxia-inducible factor (HIF) 1α and GLUT1 after 16 h medium volume change (12-well plate) in 5% or 18% oxygen incubators, with 500 µM $CoCl_2$ as positive control (n = 3 biological replicates).

(**C**) Relative RNA expression of HIF1α target genes in 3T3-L1 adipocytes cultured in either 24-well or gas-permeable Lumox plates (n = 6 biological replicates).

(**D**) Extracellular measurements of medium glucose and lactate 24 h or 48 h after medium volume change in 12-well plates (n = 3 technical replicates from n = 3 biological replicates).

(**E**) Schematic representation of the RNAseq experimental workflow. RNA extracted from 3T3-L1 adipocytes (cultured in high or low medium for 16 h in 12-well plates) or scWAT (obtained from mice kept in 10% or 21% $O_2$ for 4 weeks) were sequenced. The two sets of analysed data were then compared.

(**F**) Venn diagram showing the overlapping differentially expressed genes (p-*adj* < 0.05) from both 3T3-L1 adipocytes (high *vs* low medium) (n = 6 biological replicates) and mice scWAT (10% *vs* 21% $O_2$) (n = 10 biological replicates).

(**G**) Fold change of the 703 differentially expressed genes in 3T3-L1 adipocytes (y-axis) and mice scWAT (x-axis) from the intersection in Figure S3D, showing a 77% directional concordance.

Data are represented as mean ± SEM. *$p < 0.05$, **$p < 0.01$, ***$p < 0.001$, ****$p < 0.0001$ by paired Student’s t-tests (C), or by two-way ANOVA with Šidák correction for multiple comparisons (A, G).

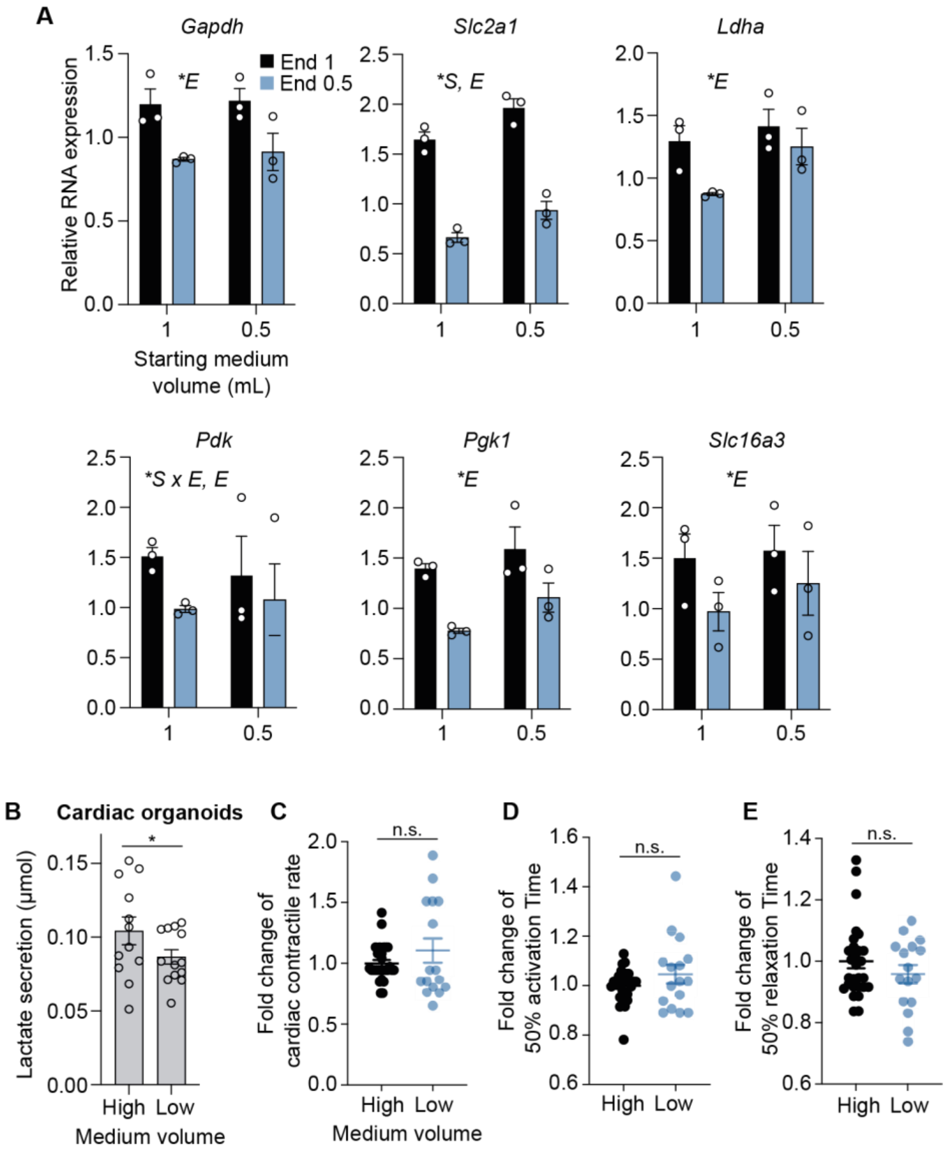


**Fig. S4.**
**Effects of low medium volumes on hPSC-derived hepatocytes and cardiac organoids**
(**A**) Relative RNA expression of HIF1α target genes in iPSC-derived hepatocytes. Cells were differentiated in either 1 mL or 0.5 mL medium (Starting medium volume), and switched to either 1 mL or 0.5 mL medium for the final 24 h (End) (n = 3 biological replicates).
(**B**) Lactate secretion by cardiac organoids after 48 h of medium volume change (n = 3-6 technical replicates from n = 3 biological replicates). High = 150 µL, Low = 50 µL.
(**C**) Contractile rate (normalised) (n = 2-17 technical replicates from n = 4 biological replicates).
(**D**) Time from 50% activation to peak (normalised) (n = 2-17 technical replicates from n = 4 biological replicates).

(**E**) Time from peak to 50% relaxation (normalised) (n = 2-17 technical replicates from n = 4 biological replicates).

All data are represented as mean ± SEM. n.s., non-significant; *$p$ < 0.05 by two-way ANOVA (A) or by unpaired two-tailed Student's t-test (B and C).
